# Neoantigen driven B cell and CD4^+^ T follicular helper cell collaboration promotes robust anti-tumor CD8^+^ T cell responses

**DOI:** 10.1101/2020.12.23.424168

**Authors:** Can Cui, Jiawei Wang, Ping-Min Chen, Kelli A. Connolly, Martina Damo, Eric Fagerberg, Shuting Chen, Stephanie C. Eisenbarth, Hongyu Zhao, Joseph Craft, Nikhil S. Joshi

**Affiliations:** Department of Immunobiology, Yale University School of Medicine, New Haven, CT 06520, USA; Department of Internal Medicine (Rheumatology, Allergy and Immunology), Yale University School of Medicine, New Haven, CT 06520, USA; Program of Computational Biology and Bioinformatics, Yale University, New Haven, CT, USA 06510, USA; Department of Lab Medicine, Yale University School of Medicine, New Haven, CT 06519, USA; Department of Biostatistics, Yale School of Public Health, New Haven, CT 06510, USA

## Abstract

CD4^+^ T follicular helper (TFH) cells provide help to B cells, which is critical for germinal center (GC) formation, but the importance of TFH-B cell interactions in cancer is unclear. We found TFH cells correlated with GC B cells and with prolonged survival of lung adenocarcinoma (LUAD) patients. To investigate further, we developed an LUAD model, in which tumor cells expressed B-cell- and T-cell-recognized neoantigens. Interactions between tumor-specific TFH and GC B cells were necessary for tumor control, as were effector CD8^+^ T cells. The latter were reduced in the absence of T cell-B cell interactions or the IL-21 receptor. IL-21 was produced primarily by TFH cells, development of which required B cells. Moreover, development of tumor-specific TFH cell-responses was also reliant upon tumors that expressed B-cell-recognized neoantigens. Thus, tumor-neoantigens themselves can control the fate decisions of tumor-specific CD4^+^ T cells by facilitating interactions with tumor-specific B cells.

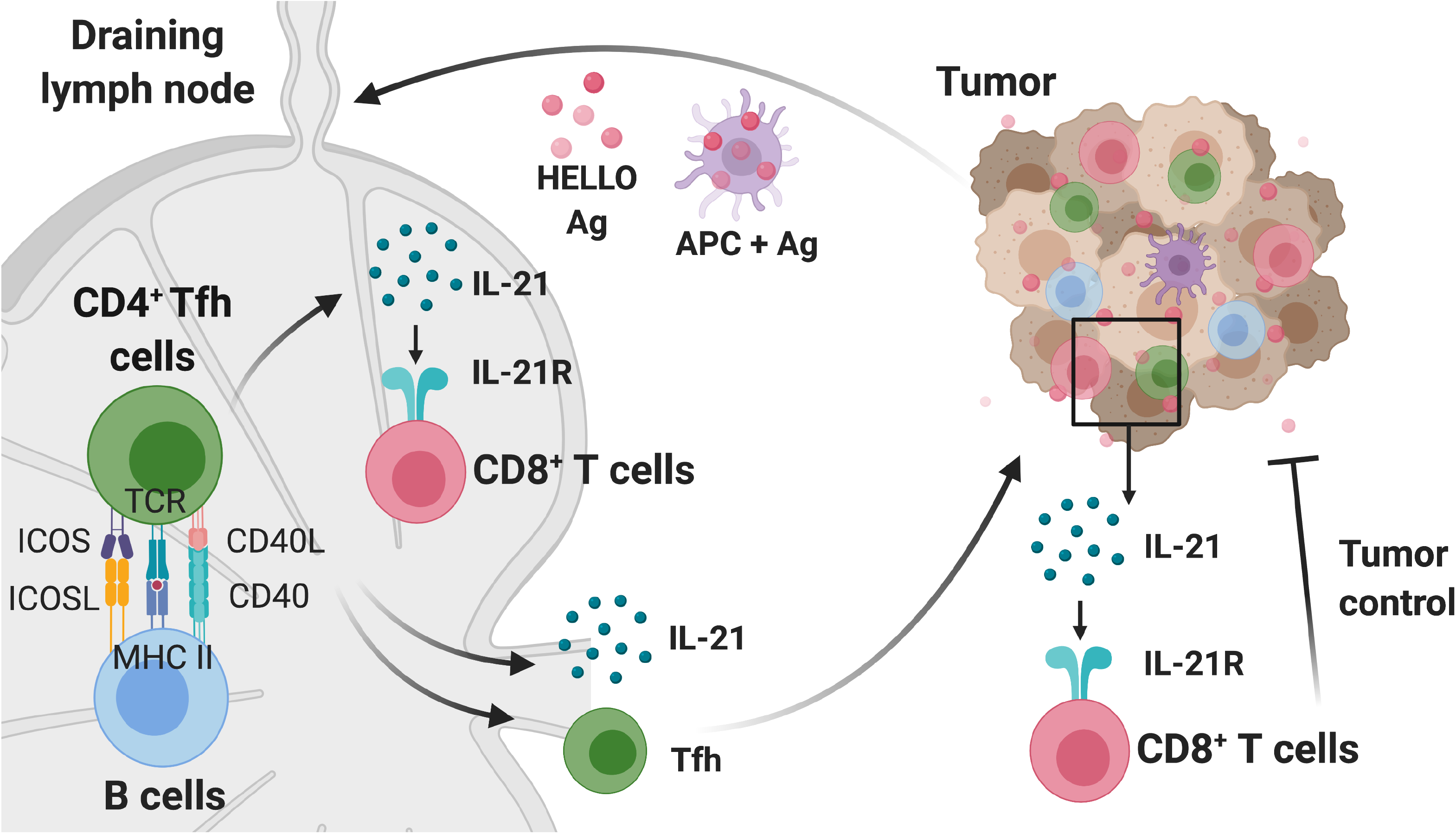

## Introduction

Lung cancer is the leading cause of cancer-related mortality worldwide, accounting for ~1.8 million deaths annually (Bray et al., 2018). The most common form of lung cancer is lung adenocarcinoma (LUAD, 40% of cases), a form of non-small cell lung cancer (NSCLC) that is most frequently associated with mutations in the KRAS proto-oncogene (30% of cases) (Schabath and Cote, 2019). Traditionally, therapeutic options for KRAS-mutant LUAD have been limited, but the successful application of immune checkpoint blockade (ICB) targeting the PD-1 pathway has changed the therapeutic landscape (Ettinger et al., 2019; Hanna et al., 2020). The success of PD-1-blockade therapy has also changed the perception that LUAD is a non-immunogenic cancer type because anti-PD-1/PD-L1 treatment potentiates the function of previously activated T cells, suggesting therapy-responsive LUAD patients had potent, ongoing anti-tumor immune responses at the time of therapy administration (Forde et al., 2018; Gettinger et al., 2018; Lizotte et al., 2016; Topalian et al., 2012). Yet, only ~20% of LUAD patients benefit from ICB (Gandhi et al., 2018; Herbst et al., 2020; Reck et al., 2016) and improving response rates will require a better understanding of the anti-tumor immune response, including how different types of tumor-infiltrating T cells interact with other immune cells and contribute to anti-tumor immunity.

A growing body of evidence has highlighted the potential importance of CD4^+^ T helper (Th) cells and B cells in cancer and immunotherapy (Alspach et al., 2019; Borst et al., 2018; Ferris et al., 2020; Helmink et al., 2020; Tran et al., 2014; Wieland et al., 2020). Much of the mechanistic research on CD4^+^ T cells has focused on Th1, Th2 and Th17 cells (Galon et al., 2013; Goc et al., 2014; Guo et al., 2018; Liu et al., 2020; Martin-Orozco et al., 2009; Oh et al., 2020), and relatively little is known about the functional role of TFH cells in cancer. In non-cancer contexts, TFH cells provide necessary help for B cell maturation and function. In a wide range of human cancers, the presence of B cells and TFH cells is correlated with prolonged survival and favorable therapeutic responses, including in NSCLC (Germain et al., 2014), breast cancer (Garaud et al., 2019; Gu-Trantien et al., 2013; Gu-Trantien et al., 2017; Hollern et al., 2019; Lu et al., 2020), melanoma (Cabrita et al., 2020; Griss et al., 2019; Helmink et al., 2020), sarcoma (Petitprez et al., 2020), head and neck cancer (Cillo et al., 2020), colorectal cancer (Bindea et al., 2013), gastric cancer (Hennequin et al., 2016) and ovarian cancer (Kroeger et al., 2016; Truxova et al., 2018). This correlation is especially strong when TFH cells and B cells are located in tumor-associated tertiary lymphoid structures (TLSs), as these structures could facilitate TFH cell-B cell interactions with subsequent effector functions in tumors (Ruffin et al., 2020; Sautes-Fridman et al., 2016; Sautes-Fridman et al., 2019; Sharonov et al., 2020). Yet, the functional importance of tumor-specific TFH cells remains uncertain, as does the importance of their interactions with B cells.

T cell-B cell interactions are necessary for the proper development and function of TFH cells and germinal center (GC) B cells in infection, vaccine responses, and autoimmune diseases (Crotty, 2019; Cyster and Allen, 2019). In canonical T-dependent immune responses, TFH cell differentiation requires interactions with two types of antigen-presenting cells. Dendritic cells (DCs) prime naïve CD4^+^ T cells by providing antigen-MHC class II complexes in concert with costimulation, including that delivered by ICOS (inducible co-stimulator) via ICOS ligand (ICOSL) on DCs. These signals, plus cytokines, initiate a transcriptional program driven by the canonical TFH cell transcription factor B-cell lymphoma 6 (Bcl-6) and a second transcription factor Achaete-scute complex homolog 2 (Ascl2), leading to downregulation of P-selectin glycoprotein ligand 1 (PSGL1), and upregulation of PD1 and the chemokine receptor CXCR5, as cells acquire the pre-TFH cell phenotype (CD4^+^ CD44^hi^ ICOS^+^ PSGL1^lo^ PD1^+^ CXCR5^+^) (Choi et al., 2013; Nurieva et al., 2008; Poholek et al., 2010). Subsequent interactions with B cells provide additional antigen-MHC II and ICOSL signals and are required for further maturation into GC-TFH cells that are capable of migrating via CXCR5 to, and functioning within, the GC. These GC-TFH cells are characterized by further upregulation of ICOS, CXCR5 and PD1 (CD44^hi^ ICOS^hi^ PSGL1^lo^ PD1^hi^ CXCR5^hi^) (Crotty, 2011). GC-TFH cells provide CD40 ligand (CD40L, CD154) and interleukin-21 (IL-21) to CD40^+^ B cells, signals necessary for the differentiation and maturation of GC B cells to memory B cells and long-lived plasma cells (Elgueta et al., 2009; Linterman et al., 2010; Zotos et al., 2010). IL-21 is the signature cytokine of TFH cells, although NKT, Th1, and Th17 cells also produce it under certain conditions (Chtanova et al., 2004; Coquet et al., 2007; Crotty, 2011; Korn et al., 2007; Nurieva et al., 2007; Suto et al., 2008; Wei et al., 2007). Thus, antigen-engaged, bidirectional TFH-B cell collaboration plays a pivotal role in the initiation and maintenance of robust humoral immune responses (Mesin et al., 2016). Yet, the importance of interactions between tumor-specific TFH cells and B cells has not been studied in cancer. Nor is it clear whether anti-tumor TFH cells provide other functions in cancer beyond B cell help.

Studies in chronic viral infection and cancer have shown that CD4^+^ T cells are critical to maintain effector CD8^+^ T cell function (Battegay et al., 1994; Borst et al., 2018; Matloubian et al., 1994; Zajac et al., 1998; Zander et al., 2019), with the latter necessary for therapeutic efficacy after ICB (Fridman et al., 2017; Huang et al., 2017; van der Leun et al., 2020). The mechanisms for maintaining functional CD8^+^ T cells in tumors remain uncertain, but one proposal involves CD4^+^ T cells providing IL-21, which drives effector CD8^+^ T cell differentiation and function, preventing exhaustion during chronic immune responses (Elsaesser et al., 2009; Frohlich et al., 2009; Ren et al., 2020; Snell et al., 2018; Xin et al., 2015; Yi et al., 2009). In line with this, CD8^+^ T cell exhaustion, a terminal differentiation process that results in diminished effector function and proliferative potential (Wherry and Kurachi, 2015), is enhanced in the absence of CD4^+^ T cells or IL-21 in chronic infection. A similar role for CD4^+^ T cell dependent IL-21 has been suggested in cancer (Zander et al., 2019), but it is not clear what the cellular source of IL-21 might be in tumors.

Given their potential importance in anti-tumor immunity, we studied the function of tumor-specific TFH cells in lung cancer and determined how their interactions with tumor-specific B cells and CD8^+^ T cells supported the anti-tumor immune responses.

## Results

### GC B cells and TFH cells correlate with favorable clinical outcomes in LUAD patients

To investigate the potential importance of GC B cells and TFH cells in human LUAD, we first analyzed RNA sequencing data from 513 LUAD samples in the TCGA database using CIBERSORT and LM22-referenced single-cell gene expression profile (GEP) matrix (Newman et al., 2015). We found that several immune cell types were enriched, including myeloid populations and CD8^+^ T cells (**Figure 1A**), while B cell lineage and CD4^+^ T cells were the first and third most abundant immune cell types (**Figure 1A**). Although the LM22 matrix does not distinguish GC B cells, the algorithm suggested that TFH cells and activated memory CD4^+^ T cells comprised a large proportion of the CD4^+^ T cells in tumors (**Figure S1A**). To more specifically delineate which types of CD4^+^ T cells and B cells were present in LUAD, we analyzed published single cell RNA sequencing data from a study that included 44 patient samples of treatment-naïve LUAD from surgical resection or endobronchial ultrasound bronchoscopy biopsy (GSE131907) (Kim et al., 2020). We cataloged 49,901 T/NK cells and 22,592 B cells from primary tumors (tLung), metastatic lymph nodes (mLN), normal lung tissues (nLung) and normal lymph nodes (nLN), based on their previous annotations. T/NK cells were grouped into 15 clusters and visualized by dimension reduction method Uniform Manifold Approximation and Projection (UMAP) (**Figure 1B**). Eight clusters were distinguished as CD4^+^ T cells, two of which displayed enrichment of TFH-signature genes. Notably, both TFH-like clusters (cluster 14 and 10) had higher expression of the TFH signature cytokine IL-21 and were more prominent in tLung than nLung (**Figure 1D**). Likewise, from nine B cell clusters, we identified one GC B cell cluster (cluster 5) and two antibody-secreting cell (ASC) clusters (cluster 4 and 6), both of which were enriched in tLung, compared with nLung (**Figure 1C and 1E;** note, cluster numbers in **Figure 1C and 1E** do not correspond with those in **Figure 1B and 1D**). These data suggested that TFH and GC B cells are enriched in human LUAD tumor tissues.

**Figure 1.**
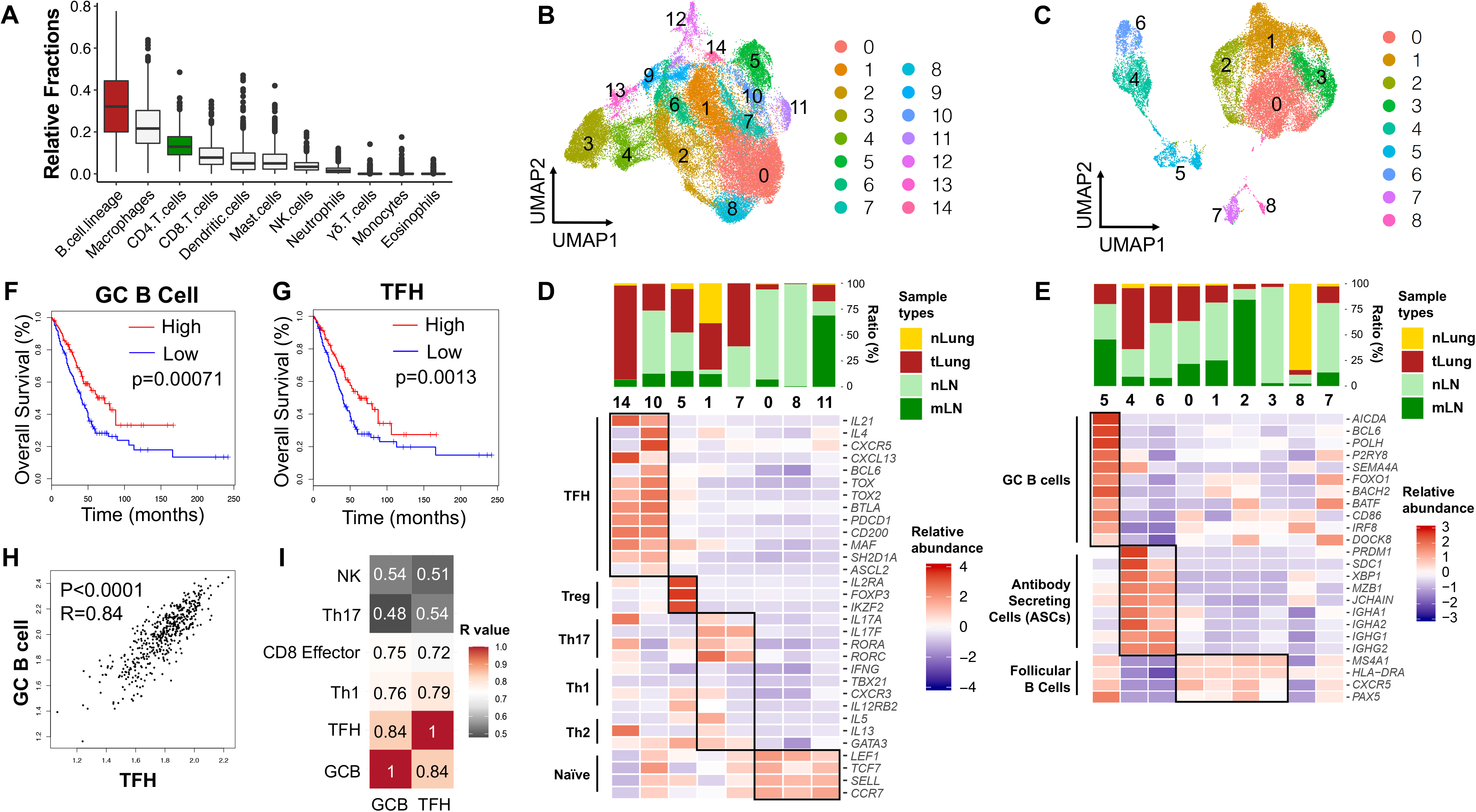
GC B cells and TFH cells correlate with favorable clinical outcomes in LUAD patients. **A.** Estimated fractions of immune cell subsets from bulk expression data of primary tumor tissues in TCGA-LUAD patient cohort (n=513). CIBERSORT algorithm and LM22 signature matrix were used for deconvolution. **B and C.** UMAP displaying T/NK cell clusters (**B**) and B cell clusters (**C**) in treatment-naïve human LUAD samples from primary sites (tLung), metastatic LNs (mLN), normal lung tissues (nLung) and normal LNs (nLN). (GSE131907, Kim *et.al.*) **D and E.** Top: tissue origins of CD4^+^ T cell clusters (**D**) and B cell clusters (**E**). Bottom: heatmap displaying relative abundance of signature genes for each cluster. Relative abundance was calculated as *Z*-scaled average of log-transformed and cell-normalized counts. **F and G.** Survival analyses based on GC B cell-signature (**F**) and TFH-signature (**G**) in lung adenocarcinoma (LUAD) patients from the TCGA (n=478). GC B cell-signature was defined as *AICDA, BATF, BACH2, BCL6, CD79A, CD79B, CD86, DOCK8, IRF4, IRF8, MYC*. TFH-signature was defined as *ASCL2, BCL6, CD4, CD200, CXCR5, IL4, IL21, IL6ST, MAF, PDCD1, SH2D1A, TOX2*. Analyses were performed with log-rank Mantel-Cox test. **H**. Correlation between the expression of GC B cell-signature and TFH-signature in TCGA-LUAD cohort. P value was calculated with two-tailed Pearson correlation test. **I**. Heatmap showing R values for the correlations among expression signatures of GC B cells, TFH, Th1, CD8 effector cells, Th17, NK cells in TCGA-LUAD cohort. Th1-signature was defined as *BHLHE40, CXCR3, CD4, IFNG, IL2, IL12RB2, STAT4, TBX21*. CD8 effector cell-signature was defined as *CD8A, CD8B, CX3CR1, GZMA, GZMB, KLRG1, PRF1, ZEB2*. Th17-signature was defined as *CD4, IL6R, IL17A, IL17F, IL23R, RORA, RORC, STAT3*. NK-signature was defined as *CD160, CD244, GNLY, GZMB, KLRC3, KLRF1, NKG2A, NKG7*. Analyses were performed with two-tailed Pearson correlation test.

To examine whether GC B cells and TFH cells associate with clinical outcomes of LUAD patients, we established gene expression signatures for GC B cells and TFH cells based on previous human studies in lymphoid tissues and cancers (Cabrita et al., 2020; Cillo et al., 2020; Milpied et al., 2018; Vella et al., 2019), and refined them based on the data in **Figures 1D – 1E** to reflect the gene expression of GC B cells and TFH cells in LUAD. Analysis of 478 LUAD cases from TCGA showed significant correlations between prolonged overall survival in LUAD patients and GC B cell- or TFH-signatures (**Figure 1F and 1G**, respectively). By contrast, Th2-, Th17- or ASC- signatures did not correlate with overall survival, and Treg-signature correlated with decreased survival (data not shown). We also identified a significant correlation between the GC B cell- and TFH-signatures (R=0.84, p < 0.0001), which suggested the co-existence of these two cell types in tumors (**Figure 1H**). GC B cell- and TFH-signatures were also correlated with Th1- and CD8^+^ effector T cell-signatures (**Figure 1I and S1B**). The correlation of TFH- and Th1-signatures was predicted as these cell types likely share a common developmental origin (Nakayamada et al., 2011; O’Shea and Paul, 2010). However, correlations between the signatures of GC B cells, TFH cells, and CD8^+^ T cells were intriguing because they suggested potential connections between these three immune cell types. By contrast, neither the GC B cell- nor TFH-signatures were correlated with the signatures for Th17 cells or NK cells. Together, these data suggested that GC B cells and TFH cells may be present in human lung cancer and contribute to anti-tumor immune responses.

### TFH cells are enriched in immunogenic murine cancer model

Next, we studied whether TFH cells were enriched in the context of murine LUAD. For this, we used a tumor cell line made from a lung adenocarcinoma genetically engineered Kras^LSL-G12D^; Trp53^fl/fl^ (KP) mouse on syngeneic C57BL/6 (B6) background (Damo et al., 2020). We subcutaneously implanted tumor cells from this KP cell line into B6 recipients. As positive controls, we implanted tumor cells from MC38 (colon cancer) and B16-F10 (melanoma) tumor cell lines, as these have been shown to elicit TFH responses *in vivo* after subcutaneous implant (Magen et al., 2019; Singh et al., 2020). We analyzed recipients for the presence of increased TFH cells in tumor-draining LNs by flow cytometry, 10 – 12 days after tumor implant. Consistent with a progressive development of CD4^+^ T cell responses, we observed ~3-fold increases in CD44^hi^ PSGL1^lo^ CD4^+^ T cells in MC38 and B16-F10 tumor-bearing mice, compared to non-tumor bearing controls, and ~11% of these PSGL1^lo^ cells further displayed upregulated CXCR5 and PD-1, indicating their development into TFH cells, likely dependent upon interactions with B cells (Weinstein et al., 2014). By contrast, in KP tumor-bearing mice we observed only a ~1.5-fold increase of CD44^hi^ PSGL1^lo^ CD4^+^ T cells, and only ~4% of these became TFH cells (**Figure S2A**). Unlike the immunogenic MC38 and B16-F10 cell lines, tumors from KP cell lines are notable in that they have low numbers of somatic mutations and are non-immunogenic (DuPage et al., 2011; McFadden et al., 2016; Yadav et al., 2014). Thus, we hypothesized that the absence of TFH cells in KP-tumor-bearing recipients likely reflected the absence of neoantigens capable of driving tumor-specific TFH cell responses.

### Engineering KP cell lines to elicit antigen-specific T cell and B cell responses

The KP tumor cell line provided a system where we could assess how engineered neoantigens would drive anti-tumor T and B cell responses. Moreover, since neither T nor B cells exhibited impact on the growth of KP cell line (**Figure S2B – S2C**; note, RAG1-knockout [RAG1 KO] mice lack T and B cells), any alterations in tumor growth would be due to T and/or B cell responses against the engineered neoantigens.

We recently published the “NINJA” model and showed that the NINJA system could be used to program neoantigen-expressing KP lung tumors (Connolly et al., 2020; Damo et al., 2020). The cDNAs encoding NINJA neoantigens contain model antigens from the lymphocytic choriomeningitis virus (LCMV) glycoprotein: GP_33-43_ and GP_61-80_, which are recognized by GP33-specific CD8^+^ (P14) and GP66-specific CD4^+^ (SMARTA) T cell receptor (TCR) transgenic T cells, respectively, separated by the FLAG peptide sequence and embedded within GFP (called GFP-GP_33-43_/FLAG/GP_61-80_).The LCMV antigens were chosen because there is a wide array of immunologic tools that are available for studying T cell responses against these antigens (*i.e.*, MHC class I and II tetramers and TCR transgenic mice). Moreover, we had previously engineered the sequence GP_33-43_/FLAG/GP_61-80_ into GFP (called GFP-GP_33-43_/FLAG/GP_61-80_ below) such that it allowed proper folding and fluorescence of GFP. Thus, we reasoned it would be an ideal starting point for a novel neoantigen that was also recognized by B cells. Hen-Egg Lysozyme (HEL) is a widely used model B cell antigen, recognized by MD4 and SW_HEL_ B cell receptor (BCR) transgenic B cells (Goodnow et al., 1988; Phan et al., 2003). Therefore, to facilitate the investigation on tumor-specific T cell and B cell responses, we created a fusion protein of **<u>HEL</u>**, **<u>L</u>**CMV GP_33-43_/FLAG/GP_61-80_, T2A, and codon-**<u>O</u>**ptimized mScarlet (**Figure 2A**). This substrate was called **<u>HELLO</u>** and generated two polypeptides: the neoantigen fusion of HEL-GP_33-43_/FLAG/GP_61-80_, and mScarlet, a bright red fluorescent protein to distinguish neoantigen-expressing tumor cells.

**Figure 2.**
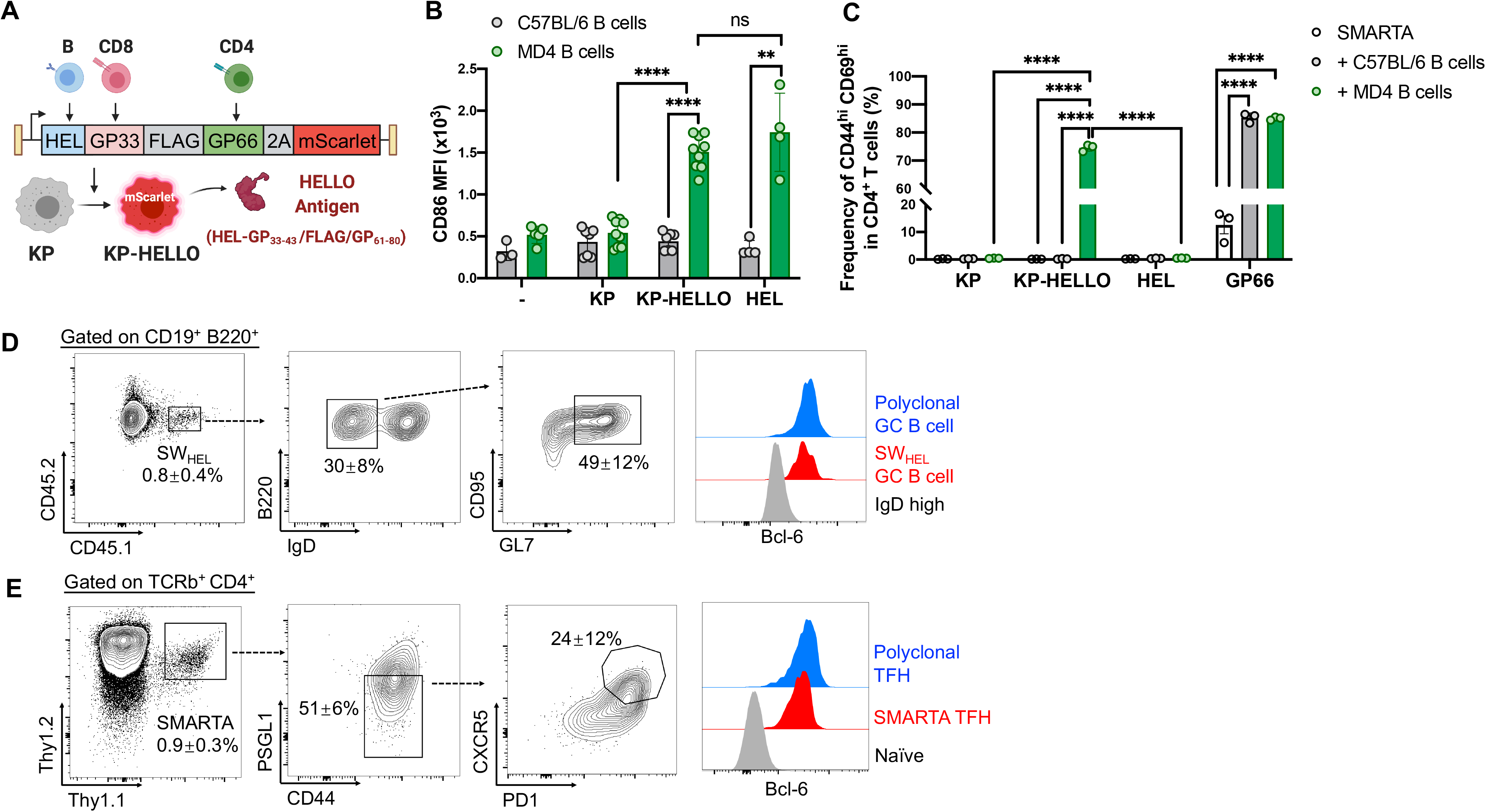
KP-HELLO tumor cells elicit tumor-specific T cell and B cell responses. **A.** Design of KP-HELLO model. **B.** Expression of CD86 on B cells (CD19^+^ B220^+^ TCRb^−^) isolated from MD4 or C57BL/6 mice, 48 hours after co-culture with the supernatant of KP, KP-HELLO, or 500ng/mL HEL. Data (mean ± SD) were pooled, from 2–5 mice/group/experiment, and representative of 2 independent experiments. **C.** Expression of CD69 and CD44 on CD4^+^ T cells isolated from SMARTA mice, 48 hours after co-cultured with B cells from MD4 or C57BL/6 mice, and the supernatant of KP, KP-HELLO, 500ng/mL HEL or 5ug/mL GP_61-80_. Data (mean ± SD) represented 2 independent experiments. **D – E,** Representative flow plots showing tumor-specific SW_HEL_ GC B cells (**D**) and SMARTA TFH cells (**E**) in tumor-draining LNs of KP-HELLO bearing mice. As shown in **Figure S4D**, CD45.2/CD45.2 Thy1.2/Thy1.2 C57BL/6 recipient, transferred with 1 × 10^5^ CD45.1/CD45.2 HEL-specific SW_HEL_ B cells and 1 × 10^5^ Thy1.1/Thy1.2 GP66-specific SMARTA CD4^+^ T cells, were subcutaneously implanted with 5 × 10^5^ KP-HELLO. Flow cytometric analyses were performed on day 10 – 12. Data (mean ± SD) represented 3 independent experiments. **B – C**, two-tailed Student’s t-test. *P < 0.05, **P < 0.01, ***P < 0.001, ****P < 0.0001.

We transduced the KP tumor cell line with HELLO-encoding lentiviral vectors, in which HELLO was the only protein product, and sorted mScarlet-positive **<u>KP-HELLO</u>** tumor cells with FACS. We placed a signal peptide sequence in the HELLO construct prior to the neoantigen fusion HEL-GP_33-43_/FLAG/GP_61-80_ so that it would be secreted by KP-HELLO cells, while mScarlet protein would remain in the cytoplasm (**Figure 2A**). To validate this, we isolated B cells from HEL-specific MD4 BCR-transgenic mice or polyclonal B cells from B6 mice that are not enriched in HEL-binding BCRs (negative control), and added either HEL protein (positive control), KP supernatant or KP-HELLO supernatant into each condition. After 48 hours of culture, flow cytometric analyses showed activation of MD4 B cells, based on a significant increase in CD86 median fluorescence intensity (MFI), in samples where MD4 cells were incubated with KP-HELLO supernatant or HEL protein (**Figure 2B**). By contrast, B cell activation was not observed in negative control conditions. Next, we tested if HEL-specific B cells could present GP_66-77_ from HEL-GP_33-43_/FLAG/GP_61-80_ in the KP-HELLO supernatant to GP66-specific CD4^+^ T cells. For this, we cultured naïve GP66-specific SMARTA CD4^+^ T cells with HEL-specific MD4 B cells, or as negative controls, SMARTA T cells alone or with polyclonal B cells from B6 mice. Into each culture condition, we added either KP supernatant, KP-HELLO supernatant, HEL protein (negative control), or GP_61-80_ peptide (positive control). As expected (Bruno et al., 2017; Kolenbrander et al., 2018; Lapointe et al., 2003), after 48 hours, there was a significant increase of CD44^hi^ CD69^hi^ activated GP66-specific SMARTA T cells in positive control samples containing GP_61-80_ peptide and with either B6 B cells or MD4 B cells, demonstrating that either cell type was capable of presenting peptide antigens to SMARTA CD4^+^ T cells via MHC class II (**Figure 2C and S3**). By contrast, only MD4 B cells were capable of activating SMARTA cells when cultured with KP-HELLO supernatant, demonstrating that a HEL-specific BCR was required for B cells to uptake, process, and present HEL-GP_33-43_/FLAG/GP_61-80_ antigen to GP66-specific CD4^+^ T cells.

### KP-HELLO tumors elicit tumor-specific TFH and GC B cell responses *in vivo*

To assess whether the introduction of HELLO neoantigens into KP tumors elicited TFH cell responses *in vivo*, we subcutaneously implanted KP or KP-HELLO tumor cells into B6 mice and analyzed endogenous CD4^+^ T cells in draining LNs using flow cytometry. On day 10 – 12, the frequency of CD44^hi^ PSGL1^lo^ polyclonal CD4^+^ T cells increased ~1.3 fold (KP-HELLO vs. KP), while there were ~4-fold increases in the frequency of these PSGL1^lo^ cells that further upregulated PD-1 and CXCR5 (**Figure S2A and S4A**), which led to a significant increase in the frequency of TFH cells (**Figure S4B)**. These data were consistent with the idea that the introduction of HELLO neoantigens was sufficient to potentiate TFH cell formation. Next, we examined whether the increase in TFH cells elicited by KP-HELLO tumors was also observed amongst tumor-specific (GP66-specific) CD4^+^ T cells by using I-A^b^/GP_66-77_ MHC class II tetramer staining. Endogenous GP66-specific CD4^+^ T cells were readily detectable in mice with KP-HELLO tumors, and moreover, ~25% of these tumor-specific CD4^+^ T cells had downregulated PSGL1, ~50% of which had upregulated PD-1 and CXCR5 (**Figure S4C**). These endogenous GP66-specific TFH cells were also marked by upregulation of Bcl-6, a transcription factor that is required for TFH cell differentiation (Johnston et al., 2009; Nurieva et al., 2009; Yu et al., 2009). Similarly, when B6 mice as recipients of transferred allotype-marked naïve HEL-specific SW_HEL_ B cells and SMARTA CD4^+^ T cells (**Figure S4D**) were challenged with KP-HELLO tumor cells, we observed ~50% of SMARTA cells were PSGL1^lo^, of which ~25% were PD1^hi^ CXCR5^hi^ Bcl-6^hi^ TFH cells (**Figure 2E**). Amongst the tumor-specific transferred SW_HEL_ B cells, ~30% were IgD^lo^, evidence of their activation, of which ~50% were GL7^+^ CD95^+^ GC-phenotype B cells, and this was accompanied by upregulated expression of Bcl-6, which is also necessary for GC B cell maturation (**Figure 2D**). Together, these data demonstrate that KP-HELLO tumors were capable of eliciting tumor-specific TFH and GC B cell responses.

### Introduction of HELLO induces TFH- and B cell-dependent tumor control

Growth of the parental KP tumors was not impacted by the presence of adaptive immune system (**Figure S2B – S2C**; also note similar growth in B6, RAG1 KO, CIITA KO [MHCII transactivator knockout] mice which have impaired CD4^+^ T cell responses, and uMT mice which lack mature B cells). By contrast, the introduction of HELLO neoantigens into KP-HELLO cells reduced tumor growth in immunocompetent hosts (B6 vs. RAG1 KO; **Figure 3A**). This was not due to a cell intrinsic growth defect as KP and KP-HELLO tumors grew similarly in RAG1 KO mice (**Figure S5A**), which suggested that the adaptive immune cells exerted impact on the growth of KP-HELLO tumors. To determine which types of adaptive immune cells were necessary, we assessed the growth of KP-HELLO tumors in CIITA KO, uMT and B6 mice treated with anti-CD8a antibodies, which efficiently depleted CD8^+^ T cells (**Figure S5B**). We observed that CD4^+^ T cells, CD8^+^ T cells and B cells were all critical for optimal control of KP-HELLO tumor growth (**Figure 3B – 3D**). Finally, we assessed whether T cell-B cell interactions were necessary for tumor control. In T-dependent immune responses, mice deficient in CD40-CD40L or ICOS-ICOSL pathways are not able to form GC B cells or GC-TFH cells, due to defects in the ability of T cells and B cells to provide mutual help (Choi et al., 2011; Ise et al., 2018; Liu et al., 2015; Nurieva et al., 2008; Renshaw et al., 1994; Zhang et al., 2020). After subcutaneous implant of KP-HELLO tumors, deficiency in ICOS or CD40L led to more rapid tumor growth, on a par with the changes observed in uMT mice (**Figure 3E – 3F**). Thus, programming KP tumors with HELLO neoantigens was sufficient to allow for the tumor control that was dependent on CD4^+^ T cells, CD8^+^ T cell, B cells, and also T cell-B cell interactions.

**Figure 3.**
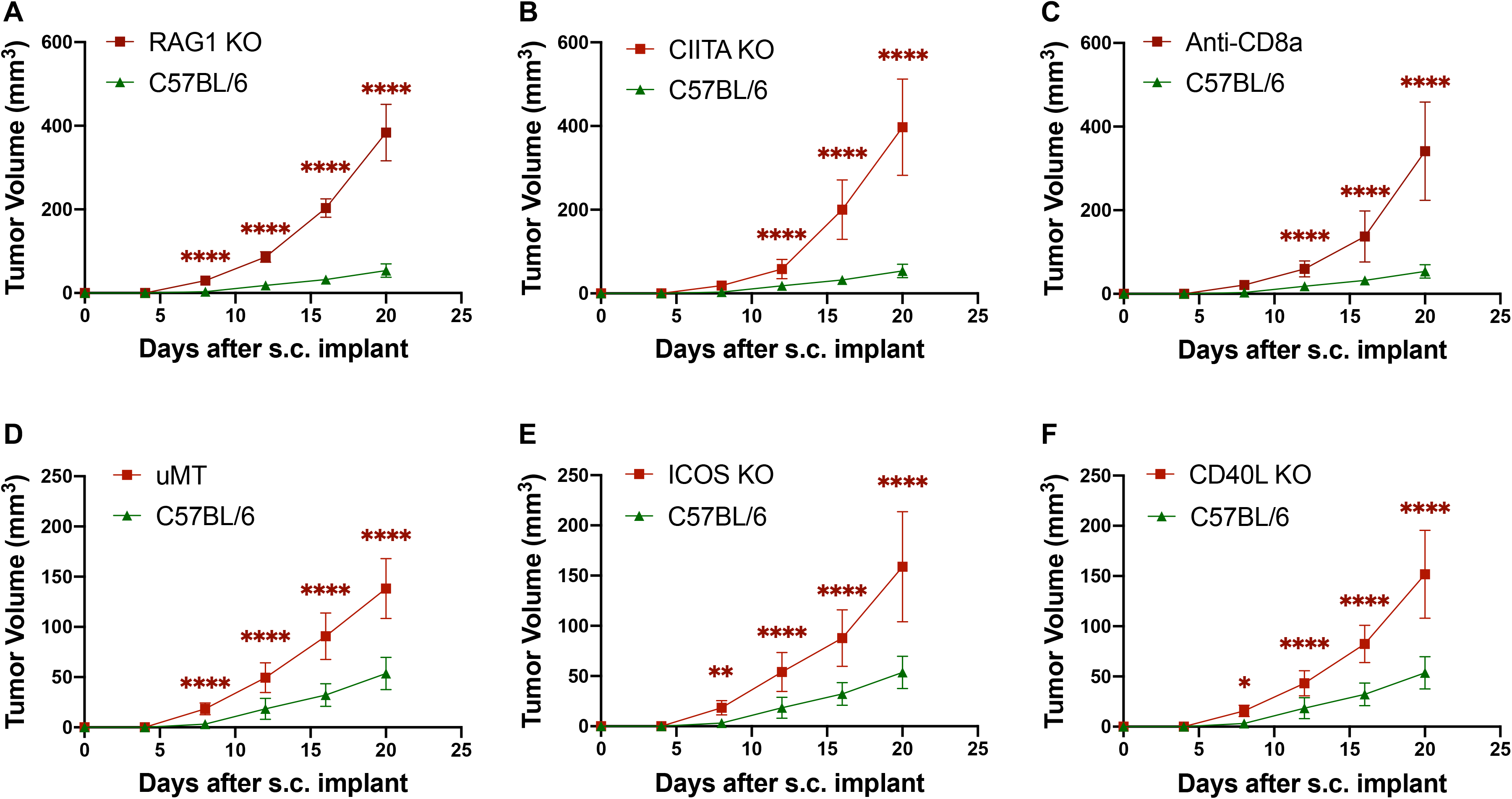
Collaborations between CD4^+^ T cells and B cells are necessary for optimal control over KP-HELLO tumors. **A – E,** Tumor growth curves: C57BL/6, RAG1 KO, CIITA KO, uMT, ICOS KO, CD40L KO mice, or C57BL/6 with CD8^+^ T cell depletion were implanted subcutaneously with 2 × 10^5^ KP-HELLO. Study end points: tumor size >1cm^3^, or signs of ulceration, infection, bleeding. Growth curves of C57BL/6 (n = 40) and uMT (n = 32), pooled from 8 independent experiments, were set as the baseline of analyses. For CD8^+^ T cell depletion, 200ug/mouse anti-CD8a antibodies (clone: 53-6.7) were administered intraperitoneally every three days, from 10 days prior to tumor initiation till study end points. Data (mean ± SD) in **A – F** were pooled, from 3–10 mice/group/experiment, and representative of at least 2 independent experiments. Analyses were performed with two-way ANOVA. *P < 0.05, **P < 0.01, ***P < 0.001, ****P < 0.0001.

### B cells are required to drive effective TFH cell responses in tumors

Given the similar increase of KP-HELLO tumor growth in uMT, CD40L KO, and ICOS KO mice, we chose to continue our studies in uMT mice, to better understand how the loss of T cell-B cell interactions impacted anti-tumor CD4^+^ and CD8^+^ T cell responses. Flow cytometric analysis of draining LNs from KP-HELLO tumor-bearing B6 and uMT mice at day 10 – 12 showed that while endogenous GP66-specific CD4^+^ T cell responses were detectable in both conditions (**Figure 4A**), GP66-specific CD44^hi^ PSGL1^lo^ PD1^hi^ CXCR5^hi^ TFH cells were absent in uMT mice (**Figure 4B and S6A**). Notably, comparing to B6 controls, uMT mice also showed ~30% and ~80% reduction in the frequency and number, respectively, of GP66-specific CD44^hi^ PSGL1^lo^ cells, comprising both TFH-precursors and TFH cells (**Figure 4C**). Moreover, in line with previous studies upon immunization or acute viral infection (Choi et al., 2011; Choi et al., 2013; Nurieva et al., 2008), Bcl-6 upregulation in CD4^+^ T cells was compromised in the absence of B cells, suggesting that interactions with B cells are necessary for the induction and/or maintenance of TFH cell differentiation in the context of tumors (**Figure S6B**).

**Figure 4.**
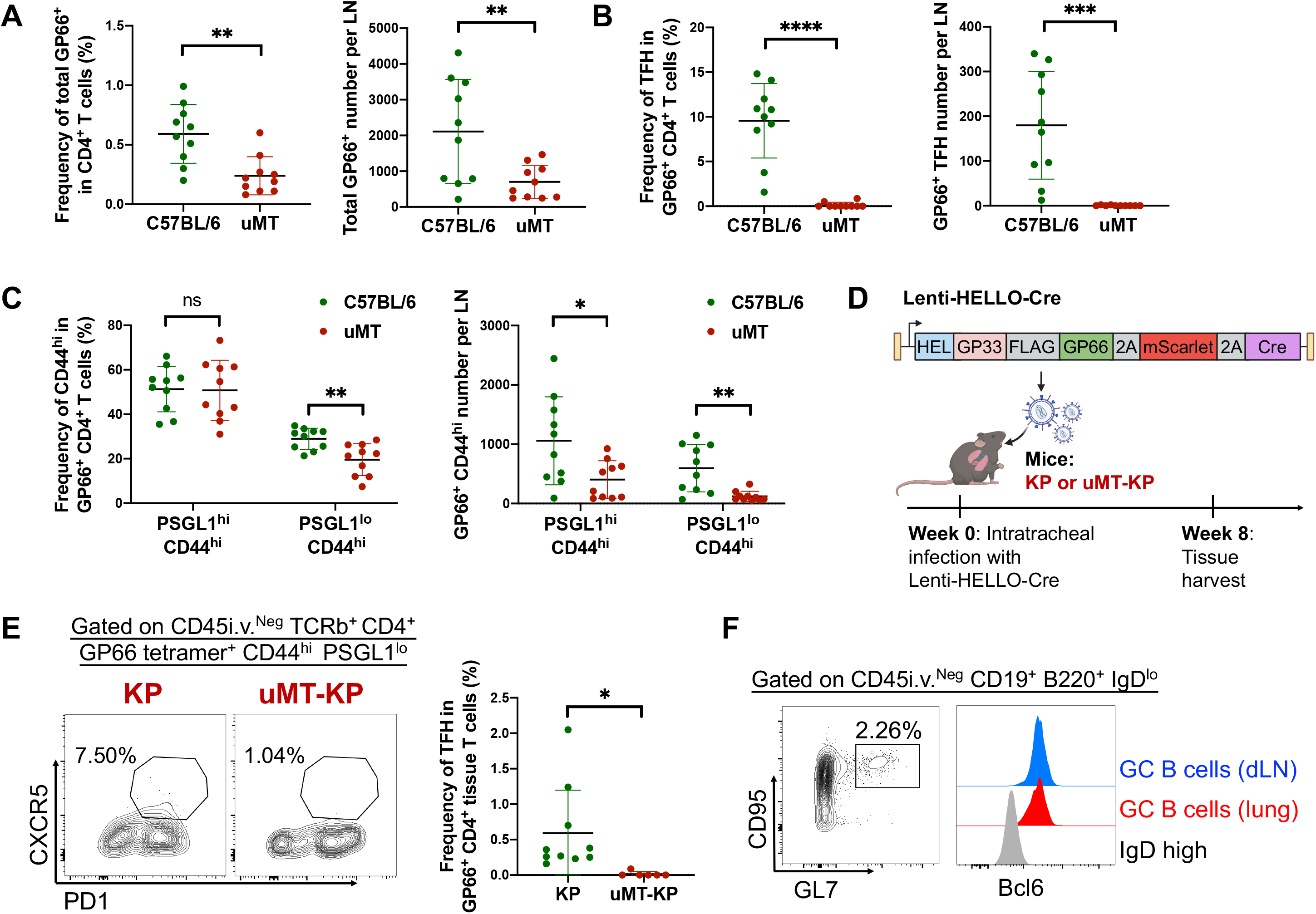
B cells are required to drive effective TFH cell responses. **A – C,** Frequency and number of I-A^b^/GP_66-77_-specific total CD4^+^ T cells (**A**), TFH cells (CD44^hi^ PSGL1^lo^ PD1^hi^ CXCR5^hi^) (**B**) and CD44^hi^ PSGL1^hi^ vs. CD44^hi^ PSGL1^lo^ CD4^+^ T cells (**C**), from draining LNs of KP-HELLO tumor-bearing C57BL/6 or uMT mice. 2 × 10^5^ KP-HELLO were implanted subcutaneously. Flow cytometric analyses were performed on day 10 – 12. **D,** Design of KP-HELLO autochthonous lung tumor model. **E – F,** KP or uMT-KP mice were intratracheally infected with 5 × 10^4^ PFU Lenti-HELLO-Cre. Tumor lung tissues and LNs were harvested at 8 weeks post-infection. **E,** Representative flow plots and frequency of I-A^b^/GP_66-77_-specific CD4^+^ TFH cells (CD44^hi^ PSGL1^lo^ PD1^hi^ CXCR5^hi^) from tumor lung tissues of KP or uMT-KP mice. Flow plots were pre-gated on CD45i.v.^Neg^ TCRb^+^ CD4^+^ GP66-tetramer^+^ CD44^hi^ PSGL1^lo^. **F,** Representative flow plots showing GC B cells in tumor lung tissues (pre-gated on CD45i.v.^Neg^ CD19^+^ B220^+^ IgD^lo^). Bcl6 expression of GC B cells in tumor lung tissue were compared to IgD^hi^ cells, and *bona fide* GC B cells in draining LNs. Data (mean ± SD) in **A – F** were pooled, from 3–10 mice/group/experiment, and representative of at least 2 independent experiments. **A – C and E**, two-tailed Student’s t-test. *P < 0.05, **P < 0.01, ***P < 0.001, ****P < 0.0001.

Next we focused on determining whether B cells and their interactions with CD4^+^ T cells are important for the development of intratumoral TFH cells in lung tumors. Lung tumors develop within lung parenchyma from previously untransformed epithelial cells and are accompanied by the development of a native tumor microenvironment (TME). The Kras^LSL-G12D^; Trp53^fl/fl^ (KP) mice recapitulate the autochthonous development of lung adenocarcinomas and the native TME, but tumors lack neoantigens within (McFadden et al., 2016). We and others have previously used lentiviral vectors to program developing lung tumors in KP mice to express T cell neoantigens (DuPage et al., 2011; Joshi et al., 2015; Pfirschke et al., 2016). Thus, we engineered Lenti-HELLO-Cre lentiviral vectors, encoding both the HELLO fusion protein and Cre recombinase, and administered them intratracheally into KP and uMT-KP mice (**Figure 4D**). In lentiviral-transduced lung epithelial cells, Lenti-HELLO-Cre led to the activation of oncogenic Kras and eliminated p53, and transformed lung cells then developed into focal lung adenocarcinomas that expressed HELLO over the course of 8 weeks. Tumors were not macroscopically visible at 8 weeks, and lungs are highly vascularized, so to distinguish lung tissue-infiltrating immune cells from circulating cells for flow cytometric analyses, we injected PE-CF594 conjugated CD45 antibodies 2-3 minutes prior to sacrifice to label circulating immune cells with CD45 (immune cells in lung tissue are CD45-PE-CF594 low called CD45i.v.^Neg^; **Figure S7A**) (Joshi et al., 2015). As expected, <1% of B, CD4^+^ and CD8^+^ T cells in non-tumor-bearing mice were in lung tissue, while ~13% of B cells, ~60% of CD4^+^ T cells, and ~ 40% of CD8^+^ T cells in HELLO tumor-bearing mice were in the lung tissue (**Figure S7B**). Consistent with our observations in human LUAD samples and subcutaneously implanted murine tumors, HELLO-expressing autochthonous tumor lung tissues also contained endogenous GP66-specific CD44^hi^ PSGL1^lo^ PD1^hi^ CXCR5^hi^ CD4^+^ TFH cells (**Figure 4E**), which expressed high levels of Bcl-6, confirming that they were mature TFH cells (**Figure S6C – S6D**). Whereas in B cell deficient uMT-KP mice with HELLO-expressing tumors, infiltrating GP66-specific TFH cells were not detected, nor was upregulation of Bcl-6 (**Figure 4E and S6C – S6D**). Concurrently, the analysis of the lung-tissue CD19^+^ B220^+^ B cells showed ~2% expressed GC B cell markers (IgD^lo^ GL7^+^ CD95^+^ Bcl6^+^) (**Figure 4F**). These data demonstrated that T cell-B cell interactions are necessary for intratumoral tumor-specific TFH cell responses in autochthonous lung adenocarcinomas.

### T cell-B cell interactions are critical for anti-tumor effector CD8^+^ T cell responses

CD8^+^ T cells were necessary for the control over KP-HELLO tumor growth (**Figure 3C**), and we speculated that this also reflected a key role for T cell-B cell interactions in driving responses by effector CD8^+^ T cells. Thus, we analyzed the impact of B cell deficiency on CD8^+^ T cells in the recipients of subcutaneously implanted KP-HELLO cells (B6 vs. uMT) and in mice with autochthonous HELLO-expressing KP lung tumors (KP vs. KP-uMT). In both cases the absence of B cells led to a significant drop in the frequency and number of endogenous tumor-infiltrating CD44^hi^ PD1^hi^ effector CD8^+^ T cells (**Figure 5A – 5B and 5D – 5E**). Specifically, CD44^hi^ PD1^hi^ cells were also marked by ~1.7-fold increase of granzyme B (**Figure 5C**). This was further validated by the reduction of tumor-infiltrating granzyme B^hi^ PD-1^hi^ CD8^+^ T cells in uMT mice (**Figure 5F**). Here, we focused on the endogenous polyclonal CD8^+^ T cell responses, because for reasons unclear and unlike what we observe in NINJA expressing tumors (Connolly et al., 2020; Damo et al., 2020), the GP33-specific CD8^+^ T cell responses against HEL-GP_33-43_/FLAG/GP_61-80_ in the context of KP-HELLO were near the limit of detection and therefore failed to meet our standards for scientific rigor.

**Figure 5.**
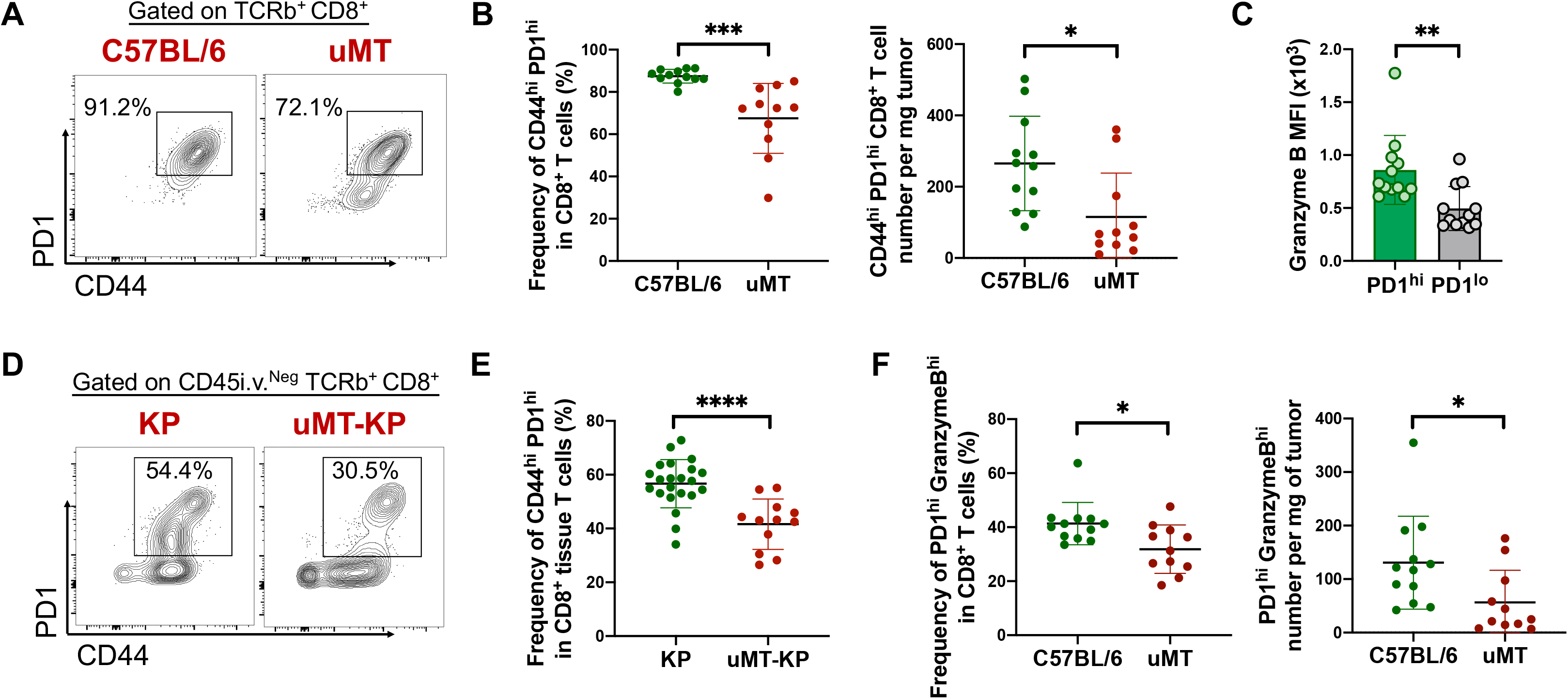
T cell-B cell interactions are critical for anti-tumor effector CD8^+^ T cell responses. **A – C and F,**2 × 10^5^ KP-HELLO were implanted subcutaneously. Tumor samples were harvested on day 15 – 16. **A – B,** Representative flow plots (**A**), frequency and number (**B**) of tumor-infiltrating CD44^hi^ PD1^hi^ CD8^+^ T cells in C57BL/6 or uMT mice with KP-HELLO tumors. **C,** Granzyme B expression on CD44^hi^ PD1^hi^ and CD44^hi^ PD1^lo^ CD8^+^ T cells in KP-HELLO tumor bearing C57BL/6 mice. **F,** Frequency and number of tumor-infiltrating PD1^hi^ Granzyme B^hi^ CD8^+^ T cells in C57BL/6 or uMT mice with KP-HELLO tumors. **D – E,** Representative flow plots (**D**) and frequency (**E**) of CD44^hi^ PD1^hi^ CD8^+^ T cells from tumor lung tissues of KP or uMT-KP mice, with autochthonous lung tumors initiated by 5 × 10^4^ PFU Lenti-HELLO-Cre administered intratracheally. Tumor lung tissues were harvested at 8 weeks post-infection. Data (mean ± SD) in **A – F** were pooled, from 3–10 mice/group/experiment, and representative of at least 2 independent experiments. **B – C and E – F**, two-tailed Student’s t-test. *P < 0.05, **P < 0.01, ***P < 0.001, ****P < 0.0001.

### IL-21 is required for robust effector CD8^+^ T cell responses in KP-HELLO tumors

Our data were consistent with the notion that interactions between B cells and CD4^+^ T cells were necessary for robust CD8^+^ effector T cell responses. We reasoned this linkage could be IL-21, as it is the signature cytokine of TFH cells (Crotty, 2011) and cognate antigen-specific B cells can promote its secretion by TFH cells (Kerfoot et al., 2011; Shulman et al., 2014). Additionally, CD4^+^ T cell-derived IL-21 has a critical role in driving effector function of CD8^+^ T cells (Snell et al., 2018; Zander et al., 2019). First, we tested whether IL-21 signals were necessary for the control over KP-HELLO tumor growth. For this, we subcutaneously implanted KP-HELLO tumor cells into B6 or IL-21 receptor knockout (IL21R KO) mice. KP-HELLO tumors grew more rapidly in IL-21R KO mice than they did in B6 mice (**Figure 6A**), consistent with a critical role for IL-21 in tumor control. To test whether the increased tumor growth was due to loss of CD8^+^ T cell function, we analyzed their infiltration into KP-HELLO tumors at day 15 – 16 and found ~60% and ~80% reduction in the frequency and number of PD1^hi^ GranzymeB^hi^ effector CD8^+^ T cells, respectively (**Figure 6B**). We also observed that tumor-infiltrating CD44^hi^ PD1^hi^ effector CD8^+^ T cells in B6 mice expressed the IL-21 receptor (**Figure 6C**). These findings were in line with the data from other groups showing that IL-21 acts on CD8^+^ T cells to drive their effector function (Snell et al., 2018; Zander et al., 2019) and confirmed the critical role of IL-21 in potentiating anti-tumor CD8^+^ T cell responses in KP-HELLO tumors.

**Figure 6.**
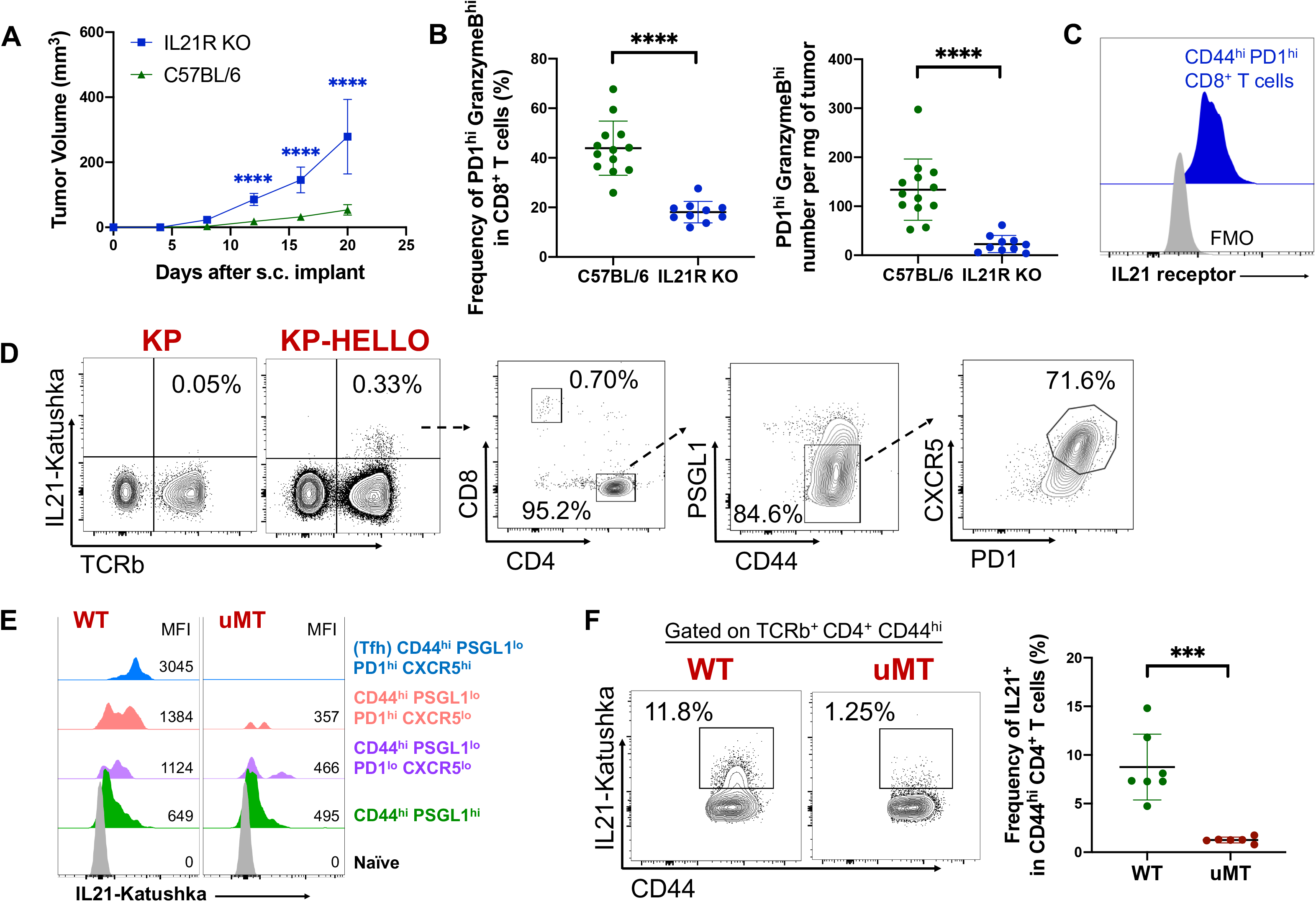
IL-21 production is dependent on B cells and is required for robust effector CD8^+^ T cell responses. **A**, Tumor growth curves: C57BL/6 and IL21R KO mice were implanted subcutaneously with 2 × 10^5^ KP-HELLO. Study end points: tumor size >1cm^3^, or signs of ulceration, infection, bleeding. Growth curve of C57BL/6 (n = 40), pooled from 8 independent experiments, was set as the baseline of analyses. **B**, Frequency and number of tumor-infiltrating PD1^hi^ Granzyme B^hi^ CD8^+^ T cells from KP-HELLO tumors in C57BL/6 or IL21R KO mice. Tumor samples were harvested on day 15 – 16. **C**, Expression of IL21 receptor on CD44^hi^ PD1^hi^ CD8^+^ T cells from KP-HELLO tumors. **D**, Representative flow plots showing IL21-expressing TFH cells in draining LNs of KP or KP-HELLO tumor-bearing mice. Flow cytometric analyses were performed on day 10 – 12. **E**, Histogram displaying the expression of IL21 from naïve, I-A^b^/GP_66-77_-specific CD44^hi^ PSGL1^hi^, CD44^hi^ PSGL1^lo^ PD1^lo^ CXCR5^lo^, CD44^hi^ PSGL1^lo^ PD1^hi^ CXCR5^lo^ and TFH cells (CD44^hi^ PSGL1^lo^ PD1^hi^ CXCR5^hi^) in draining LNs of KP-HELLO tumor-bearing WT or uMT mice. Gating strategy was shown in **Figure S6A**. **F**, Representative flow plots and summary data showing the frequency of IL21 producers in CD4^+^ CD44^hi^ T cells, from draining LNs of KP-HELLO tumor-bearing WT or uMT mice. Data (mean ± SD) in **A – F** were pooled, from 3 – 10 mice/group/experiment, and representative of at least 2 independent experiments. **A,** two-way ANOVA. **B and F**, two-tailed Student’s t-test. *P < 0.05, **P < 0.01, ***P < 0.001, ****P < 0.0001.

### IL-21 is produced primarily by TFH cells

To examine which cell types express IL-21, we subcutaneously implanted KP-HELLO tumor cells into *Il21^Kat/Kat^* reporter mice (Weinstein et al., 2016). Increased IL-21 expression was detected from T cells in KP-HELLO tumor-bearing hosts, but not in hosts with KP tumors, demonstrating that IL-21 expression was dependent on the presence of HELLO neoantigens (**Figure 6D**). Moreover, >95% of the IL-21 was expressed by CD4^+^ T cells, most of which was expressed by CD4^+^ CD44^hi^ PSGL1^lo^ PD1^hi^ CXCR5^hi^ TFH cells (**Figure 6D**). We also observed that GP66-specific TFH cells expressed IL-21, and that IL-21 expression progressively increased over different stages of TFH cell differentiation (**Figure 6E**; gating strategy in **Figure S6A**). This key finding was not restricted to KP-HELLO tumors, as IL-21 was expressed primarily by TFH cells and upregulated progressively over TFH differentiation in mice with MC38 and B16-F10 tumors (**Figure S8A – S8C**).

### B cells are necessary for IL-21 production by TFH cells

To close the loop between diminished tumor-specific TFH responses in the absence of T cell-B cell interactions, and reduced effector CD8^+^ T cells in the absence of B cells or IL-21R, we next addressed whether IL-21 expression in T cells was B-cell-dependent. For this, we bred IL-21 reporter mice to uMT mice and compared IL-21 expression in total CD44^hi^ CD4^+^ T cells between KP-HELLO tumor-bearing WT and B-cell-deficient mice. Here, we focused on CD44^hi^ CD4^+^ T cells because those comprised the majority (>90%) of the IL-21-expressing cell population (**Figure 6D**). In the absence of B cells, we observed ~85% reduction in the frequency of IL-21 expressing CD44^hi^ CD4^+^ T cells (**Figure 6F**). Analysis of GP66-specific CD4^+^ T cells in these uMT mice also showed that IL-21 expression was compromised at each stage of TFH cell differentiation, demonstrating the key role that B cells play in IL-21 expression (**Figure 6E**). Together, these data suggested that the development and/or maintenance of IL-21-expressing TFH cells requires T cell-B cell interactions.

### Tumor neoantigens regulate the cell fate decisions of anti-tumor CD4+ T cells

The experiments to this point left open the possibility that B cells act in a non-antigen driven fashion to promote the development of tumor-specific TFH cells. To formally examine whether tumor-specific B cells were required, we performed a rescue experiment by transferring 1 × 10^6^ naïve SW_HEL_ or polyclonal B6 B cells into uMT mice and challenging them with KP-HELLO tumors. Adoptive transfer of the SW_HEL_ B cells, but not B6 B cells, restored the ability of the immune response to control KP-HELLO tumors back to amounts similar to what was observed in C57BL/6 control mice (**Figure 7A**). This demonstrated that tumor-antigen-specific B cells were required to drive anti-tumor immune responses in the KP-HELLO model.

**Figure 7.**
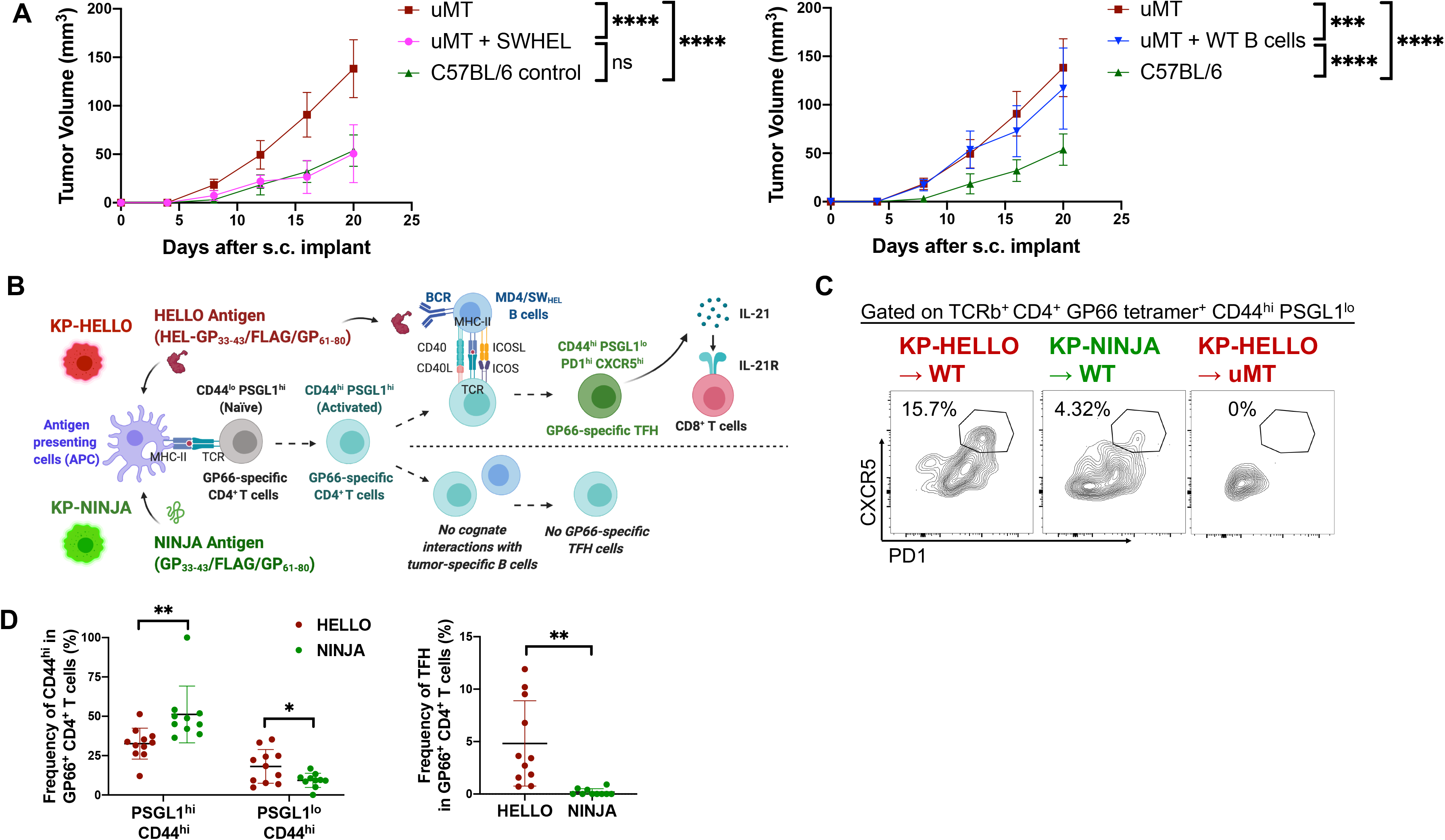
Tumor neoantigens regulate the cell fate decisions of anti-tumor CD4^+^ T cells. **A**, 1 × 10^6^ SW_HEL_ B cells or 1 × 10^6^ WT B cells were adoptively transferred to uMT recipients prior to tumor initiation. 2 × 10^5^ KP-HELLO were implanted subcutaneously. Study end points: tumor size >1cm^3^, or signs of ulceration, infection, bleeding. Growth curves of C57BL/6 (n = 40) and uMT (n = 32), pooled from 8 independent experiments, were set as the baseline of analyses. **B**, Model for **Figure 7C – 7D**. **C and D,** Representative flow plots (**C**) and frequency (**D**) of I-A^b^/GP_66-77_-specific CD4^+^ TFH cells (CD44^hi^ PSGL1^lo^ PD1^hi^ CXCR5^hi^), and frequency of I-A^b^/GP_66-77_-specific CD44^hi^ PSGL1^hi^ vs. CD44^hi^ PSGL1^lo^ CD4^+^ T cells (**D**) from tumor draining LNs. WT or uMT mice were implanted subcutaneously with 5 × 10^5^ KP-HELLO or KP-NINJA. Flow cytometric analyses were performed on day 10. Data (mean ± SD) in **A – D** were pooled, from 3 – 10 mice/group/experiment, and representative of at least 2 independent experiments**. A,** two-way ANOVA. **D,** two-tailed Student’s t-test. *P < 0.05, **P < 0.01, ***P < 0.001, ****P < 0.0001.

The HEL-GP_33-43_/FLAG/GP_61-80_ neoantigens contain epitopes that are recognized by both tumor-specific B cells and tumor-specific CD4^+^ T cells, but it is likely that neoantigen-containing proteins in human or murine tumors will fall into two categories: those that are recognized by B cells and those that are not. As antigen-recognition by B cells was critical for the uptake, processing and presentation of HEL-GP_33-43_/FLAG/GP_61-80_ neoantigens to GP66-specific CD4^+^ T cells (**Figure 2C**), we next examined whether tumor-neoantigens themselves had the ability to dictate the fate decisions of CD4^+^ T cells. To test this directly, we took advantage of KP-NINJA lung tumor cell lines as they expressed the GP_33-43_/FLAG/GP_61-80_ neoantigens as part of a flexible loop region between β-strands 7 and 8 of GFP (Damo et al., 2020). Critically, the GFP-GP_33-43_/FLAG/GP_61-80_ retains fluorescence and is recognized by conformation-specific antibodies, and thus would be unlikely to elicit GFP-specific T or B cell responses in recipient mice tolerant to GFP – for this, we used *Il4^GFP^ [4get]* mice as GFP tolerant recipients (Weinstein et al., 2016). Yet, GFP-tolerant recipients were still able to mount GP66-specific CD4^+^ T cell responses against the neoantigen-containing GFP-GP_33-43_/FLAG/GP_61-80_ in NINJA (Damo et al., 2020). Therefore, KP-NINJA and KP-HELLO systems allowed us to set up an experiment to directly compare the differentiation of endogenous tumor-specific CD4^+^ T cells under conditions where tumors expressed neoantigens with a shared CD4^+^ T cell epitope but with different capabilities to elicit B cell responses (**Figure 7B**). We subcutaneously implanted 5 × 10^5^ KP-NINJA or KP-HELLO tumors into *Il4^GFP^* mice and analyzed GP66-specific CD4^+^ T cells for their expression of TFH markers. Critically, the frequency of tumor-specific CD4^+^ CD44^hi^ PSGL1^lo^ PD1^hi^ CXCR5^hi^ TFH cells decreased ~96% in mice with KP-NINJA tumors when compared to KP-HELLO (**Figure 7C – 7D**). Moreover, although there was a 50% reduction in the frequency of GP66-specific CD44^hi^ PSGL1^lo^ population, the most prominent decrease was contributed by PD1^hi^ CXCR5^hi^ cells. This was in line with previous findings that B-cell-recognized antigens and the cognate-interactions with antigen-specific B cells were essential to drive PD1^hi^ CXCR5^hi^ TFH cell maturation in non-tumor contexts (Kerfoot et al., 2011; Shulman et al., 2014). Concomitantly, CD44^hi^ PSGL1^hi^ cells increased by ~1.6 fold (**Figure 7D**). Together these data are consistent with the hypothesis that tumor neoantigens control the developmental fates of tumor-specific CD4^+^ T cells based on whether they can also trigger responses by tumor-specific B cells.

## Discussion

Previous work has highlighted the individual roles for T cells and B cells in cancer, but less was known about the interactions between T and B cells and how they regulate anti-tumor immunity. We have identified a novel mechanism by which T cell-B cell collaboration promoted the development of TFH and GC B cells and contributed to LUAD tumor control. Tumor-specific B cells both presented tumor antigens and furnished signals that were necessary for the differentiation of IL-21-producing TFH cells. IL-21, in turn, potentiated effector functions of CD8^+^ T cells, resulting in tumor growth suppression. Yet, not all tumor neoantigens could trigger this B cell-TFH cell-IL21 pathway, as neoantigens had to be recognized by both tumor-specific B cells and tumor-specific CD4^+^ T cells. This finding underscored a novel mechanism by which tumor neoantigens could guide CD4^+^ T cell differentiation. In support of this pathway being relevant in LUAD patients, we found that human LUAD tumors contained both GC B cells and IL-21-producing TFH cells, and that there were strong correlations among signatures of GC B cells, TFH cells, and CD8^+^ T cells. Moreover, these signatures correlated with better clinical outcome for patients. Together, our findings demonstrated the pivotal role of tumor neoantigens in driving T cell-B cell collaborations, which were critical to promote anti-tumor immune responses.

We found tumor-infiltrating TFH cells and GC B cells in our analyses of human LUAD and our autochthonous KP model. The former is in line with the observations that NSCLC and other cancer types contain tumor-associated TFH-like cells with an enrichment of a TFH signature (i.e., *PDCD1, ICOS, CD200, MAF and IL21*) (Bindea et al., 2013; Cillo et al., 2020; Gu-Trantien et al., 2013; Gu-Trantien et al., 2017; Guo et al., 2018; Li et al., 2019; Satpathy et al., 2019; Thommen et al., 2018). In human studies including autoimmune diseases, TFH-like cells have been shown to express CXCL13, the ligand of CXCR5, which was associated with CXCR5^+^ B cell infiltration and tertiary lymphoid structure (TLS) formation (Bocharnikov et al., 2019; Li et al., 2015; Manzo et al., 2008; Morita et al., 2011; Rao et al., 2017). These CXCL13^+^ TFH-like cells have recently been termed peripheral helper CD4 T (Tph) cells and, under certain settings, exhibited low expression of CXCR5, reflecting potential extrafollicular location. Consistent with these findings, we observed two TFH-like clusters with high *IL-21* expression in human LUAD (**Figure 1D**): cluster 10, which was more TFH-like and enriched in both primary tumor tissues and normal LNs; and cluster 14, which was more Tph-like and primarily distributed in only tumor tissues. Due to technical reasons, we were unable to determine whether some of IL-21-producing TFH cells in our murine tumor model also expressed CXCL13. Further work will be required to understand if heterogeneity exists amongst GP66-specific TFH cells in the KP-HELLO and KP-NINJA models and to determine their significance in anti-tumor immunity.

Several studies have focused on the critical role of quiescent, stem-like CXCR5^+^ TCF1^+^ CD8^+^ T cells (TSL) in chronic immune responses. TSL cells are important for sustaining T cell responses during chronic infection and for therapeutic responses after PD-1 blockade (He et al., 2016; Im et al., 2016; Leong et al., 2016; Siddiqui et al., 2019; Utzschneider et al., 2016). Moreover, in chronic LCMV infection, CD8^+^ T cell-intrinsic IL-21R expression promotes TSL cells to differentiation into granzyme B^hi^ effector T cells (Snell et al., 2018; Zander et al., 2019). Here, adoptive transfer of *in vitro* generated IL-21-expressing CD4^+^ T cells improved CD8^+^ T cell function and led to a reduction in tumor growth. CD4^+^ T cells and IL-21 played a crucial role in maintaining virus-specific CD8^+^ T cell responses in chronic infection, and the kinetics of T cell exhaustion were faster in their absence (Elsaesser et al., 2009; Frohlich et al., 2009; Ren et al., 2020; Xin et al., 2015; Yi et al., 2009). Thus, we believe that TFH cells are a key physiologic source of IL-21 for effector CD8^+^ T cells, and that this is one mechanism for how TFH cells promote better outcomes for LUAD patients.

A critical question arising from our work is how and where TFH cells provide IL-21 to CD8^+^ T cells. During chronic LCMV infection, splenic TSL cells expressed CXCR5 and ~30% were in B cell follicles while ~50-60% in T cell zones (He et al., 2016; Im et al., 2016; Leong et al., 2016). Moreover, stem-like CXCR5^+^ CD8^+^ T cells were also enriched in tumors from NSCLC patients, and this correlated with tumor infiltration of CXCR5^+^ TFH cells and CD19^+^ B cells (Brummelman et al., 2018). Tumor-associated TLSs were observed in many types of cancers, including up to 70% of NSCLC patients, and their presence correlated with favorable clinical outcomes (Dieu-Nosjean et al., 2008; Germain et al., 2014; Goc et al., 2014; Sautes-Fridman et al., 2016; Sautes-Fridman et al., 2019). Moreover, TLSs with mature GCs were further associated with more clinical benefits for lung cancer and HPV^+^ HNSCC patients (Ruffin et al., 2020; Silina et al., 2018). Thus, the follicular structures of TLSs could support interactions between IL-21-producing TFH cells and CXCR5^+^ CD8^+^ T cells in tumors. We previously described that neoantigen-expressing KP lung tumors were associated with TLSs but did not observe the presence of GCs [(Joshi et al., 2015) and data not shown]. Further studies will be necessary to determine if the introduction of HELLO neoantigens into KP lung tumors could promote tumor-associated TLSs with GCs, and whether B cell follicles in TLSs or draining LNs are the sites of tumor-specific TFH cell-TSL cell interactions.

Studies on neoantigen immunogenicity generally focus on neoantigen abundance, MHC binding and T cell recognition as key metrics of identifying candidate neoantigens for therapeutic efforts (Schumacher and Hacohen, 2016; Schumacher et al., 2019; Wells et al., 2020); however, our findings raise the possibility that other aspects of neoantigens should also be considered. These include the assessment of mutation sites and whether they would result in structural alterations that are able to change BCR binding and B cell recognition. B-cell-recognized neoantigens were unique in that they were necessary to potentiate TFH cell differentiation pathways, which provided non-redundant help for anti-tumor CD8^+^ T cell responses via IL-21 in our model. This was in line with multiple studies showing that the presence of cognate B cells, so as cognate-B-cell-recognized antigens bearing a CD4^+^ T cell epitope, were required for IL-21 production by PD1^hi^ CXCR5^hi^ TFH cells (Kerfoot et al., 2011; Shulman et al., 2014). Moreover, the principle of intermolecular help, in which antigen-engaged B cells are able to internalize and process whole antigenic proteins, and subsequently present different MHCII epitopes derived to CD4^+^ T cells, ensures that rare, cognate interactions between tumor-specific B and CD4^+^ T cells are enabled and augmented by B-cell-recognized neoantigens (Milich et al., 1987; Mitchison, 1971a, b; Russell and Liew, 1979; Sette et al., 2008). MHC class II restricted neoantigens, recognized by CD4^+^ T cells, have been shown indispensable for successful anti-tumor responses (Alspach et al., 2019; Kreiter et al., 2015; Linnemann et al., 2015), but less is known about how individual neoantigens, or their structures, could impact the fate decisions by anti-tumor CD4^+^ T cells. We found intermolecular help was necessary for TFH cell development, and concurrently, co-existence of GC B cells and TFH cells in many human cancers likely indicated the presence of B-cell-recognized conjugate neoantigens bearing T cell epitopes. Thus, tumors not only shape T cell responses via their production of chemokines, cytokines, and innate immune signaling modulators (*i.e.*, STING/ RIG-I), but also via structural features of their neoantigens. Therefore, gaining a better understanding of how various features of neoantigens drive different CD4^+^ T cell fates could help the designing of novel therapies to leverage a wider variety of potential functions that anti-tumor CD4^+^ T cells could provide. Our work provided a novel and easily applicable platform to investigate different aspects of B and T cell neoantigens, such as secreted/membrane-bound/intracellular features, and BCR binding affinity, of which available tools have already been established from many studies (Burnett et al., 2018; Paus et al., 2006). In addition, the Gibson assembly-based modular assembly platform developed by us allows for a fast adaptation of neoantigen programming within a short period of time (Akama-Garren et al., 2016).

Given the pivotal functional role of T cell-B cell interactions in controlling tumors, demonstrated by our work and others, the B cell-TFH cell-IL21 axis could provide a novel entry point for therapeutic manipulation. Preclinical studies and phase I clinical trials of personalized neoantigen vaccines, utilizing MHC class I binding epitopes, have successfully demonstrated the induction of *de novo* neoantigen-specific CD4^+^ and CD8^+^ T cell responses in patients with NSCLC, bladder cancer, melanoma and glioblastoma (Keskin et al., 2019; Ott et al., 2017; Ott et al., 2020). Here, one possibility could be to modify these personalized neoantigen vaccines to include epitopes recognized by both B and T cells. We predict that this would boost responses by neoantigen-specific TFH cells, facilitate their IL-21 production, and promote anti-tumor responses by effector CD8^+^ T cells. Alternatively, one study found that B cell-based neoantigen vaccines could promote checkpoint blockade efficacy by activating autologous CD8^+^ T cells in animal models of glioblastoma (Lee-Chang et al., 2021). Beyond vaccines, agonist antibodies or cytokines targeting the B cell-TFH cell-IL21 axis could also provide therapeutic benefits. Anti-ICOS agonist antibody (GSK3359609) is currently in phase II/III clinical trials in combination with PD-1 or PD-L1 blockade. Moreover, combining engineered IL-21 with existing immunotherapies can elicit robust anti-tumor immune responses (Deng et al., 2020; Singh et al., 2011; Sondergaard and Skak, 2009; Wang et al., 2019; Štach et al., 2018). Taken together, these data suggested that therapeutic designs leveraging the B cell-TFH cell-IL21 axis could advance existing immunotherapies and lead to improved response rates for patients with lung and other cancers.

## METHOD DETAILS

### Mice

All mice were housed in pathogen-free conditions at Yale School of Medicine (New Haven, CT). CD45.2 WT C57BL/6 mice, and CD45.1 allotype-marked mice were purchased from the Jackson Laboratory and Charles River Laboratory. uMT, ICOS KO, IL21R KO, and MD4 mice were obtained from the Jackson Laboratory. Class II major histocompatibility complex (MHCII) transactivator (CIITA) KO and CD40 ligand KO mice were provided by Stephanie Eisenbarth (Yale University), and RAG1 KO by David Schatz (Yale University). SMARTA, P14 and KP mice were described previously (Damo et al., 2020; DuPage et al., 2011; Joshi et al., 2015). SW_HEL_ mice were provided by Robert Brink (Garvan Institute of Medical Research). *Il21^Kat/Kat^* and *Il4^GFP^ [4get]* reporter was described previously (Weinstein et al., 2016). uMT-*IL21 ^Kat/Kat^* reporter and uMT-KP were bred in-house. All mice were used in accordance with protocols approved by the Institutional Animal Care and Use Committees of Yale University.

### Generation of KP-HELLO cell line

The KP cell line had been described previously (Damo et al., 2020) and was derived from an Ad-CRE-infected Kras^G12D^, p53^fl/fl^ mouse (DuPage et al., 2009). KP was transduced with HELLO Lentiviral vector (LV). The LV backbone for HELLO has been described previously (Akama-Garren et al., 2016). HELLO-LV contains HELLO driven by phosphoglycerate kinase (pGK) promoter. HELLO was generated as a protein fusion of codon-optimized hen egg lysozyme (HEL), LCMVgp_33-43_ (KAVYNFATCGI), LCMVgp_61-80_ (GLNGPDIYKGVYQFKSVEFD), and 2A peptide-linked mScarlet. KP cells were transduced with HELLO-LV produced by co-transfection of 293FS* cells with HELLO, PSPax2 (gag/pol) and VSV-G (vesicular stomatitis virus glycoprotein) vectors as described previously (Damo et al., 2020). Transduced LGKP cells were FACS-sorted, yielding a pure mScarlet population of KP-HELLO cells. 25 frozen aliquots of KP-HELLO were generated after 2 passages and were named as “G0”. Based on 1 aliquot of “G0”, 25 frozen aliquots of “G1” were generated after 2 passages, and so as “G2” based on “G1”. For each experiment, one fresh aliquot of “G2” was thawed and used within 2 passages. After 6-7 passages described above, >99% of KP-HELLO “G2” cells were still mScarlet positive with flow cytometric analysis, which suggested the stable expression of HELLO protein in KP-HELLO cell line.

### Lentiviral vector production and administration

“Lenti-HELLO-Cre” plasmids were generated by cloning Cre into HELLO-LV. Lentiviral vectors were produced by transfecting 293FS* cells with Lenti-HELLO-Cre, PSPax2 and VSV-G vectors at a 4:3:1 ratio, of which supernatant was harvested at 100,000g for 2 hours (20°C), after 32-hour incubation at 37°C. LV titer was quantified by transducing GreenGo cell line (10^5^ cells/well) in serial dilutions to 1mL of final volume and calculated with the formula: (% fluorescent cells × 10^5^ × Dilution Factor)/100 (pfu/mL). For autochthonous KP-HELLO tumor initiation, 5 × 10^4^ pfu Lenti-HELLO-Cre were administered intratracheally to KP mice.

### *In vitro* T cell and B cell co-culture

KP and KP-HELLO cells had been cultured at 37°C and 5% CO2 in complete RPMI-1640 (10% HI-FBS + 1% Pen/Strep + 1x L-Glut + 55μM β-mercaptoethanol) for 2 days. Supernatant of KP and KP-HELLO were filtered with 0.45μm filters. Naïve CD4^+^ T cells and B cells were separated from spleens by negative selection with EasySep Mouse CD4^+^ T Cell Isolation Kit and Mouse B Cell Isolation Kit, respectively (STEMCELL technologies), according to the manufacturer’s protocol. 1 × 10^6^/mL B cells were plated with CD4^+^ T cells in a ratio of B:T=1:2, and co-cultured with the supernatant of KP or KP-HELLO, or complete RPMI-1640 supplemented with LCMV GP_61-80_ peptide GLKGPDIYKGVYQFKSVEFD (5μg/mL, Anaspec) or HEL (0.5μg/mL, Sigma-Aldrich). 10ng/mL BAFF were added in all the co-culture experimental conditions. Flow cytometric analysis were performed after 48 hours of co-culture.

### Antibody depletion, adoptive transfer, and tumor experiments

For depletion of CD8^+^ T Cells, 200ug/mouse anti-CD8a antibodies (clone: 53-6.7; InVivoMab) were diluted in PBS, and injected intraperitoneally every three days, 10 days prior to tumor initiation until the end point of experiments. For adoptive transfer, naïve CD8^+^ T cells, CD4^+^ T cells and B cells were separated from spleens by negative selection with EasySep Mouse CD8^+^ T Cell Isolation Kit, Mouse CD4^+^ T Cell Isolation Kit and Mouse B Cell Isolation Kit, respectively (STEMCELL technologies). For co-transfer experiments, CD45.1/CD45.2 or Thy1.1/Thy1.2 allotype-marked P14 (5 × 10^5^ /mouse), SMARTA (1 × 10^5^ /mouse) and SW_HEL_ (1 × 10^5^ /mouse) cells were co-transferred into naïve CD45.2/CD45.2 Thy1.2/Thy1.2 mice that were implanted with tumor cell lines one day later. For rescue experiments, 1 × 10^6^ SW_HEL_ cells or WT C57BL/6 B cells were transferred. KP, KP-HELLO, MC38 and B16-F10 cells were cultured in complete DMEM (4.5g/L D-glucose + L-glutamine + 110mg/L sodium pyruvate + 10% HI-FBS + 1% Pen/Strep). For tumor growth measurements and FACS analysis of endogenous antigen-specific immune cells, mice were subcutaneously (s.c.) implanted with 2 × 10^5^ KP, KP-HELLO, or 5 × 10^5^ MC38, B16-F10 cells. For experiments with co-transfer of CD8^+^ T cells, CD4^+^ T cells and B cells, mice were s.c. implanted with 5 × 10^5^ KP-HELLO cells. Tumor volume was measured as [(Width^2^ × Length)/2], every four days with a digital caliper. Growth curves of C57BL/6 (n = 40) and uMT (n = 32) mice that were challenged with 2 × 10^5^ KP-HELLO cells, pooled from 8 independent experiments, were set as the baseline of analyses. Mice with ≧ 1cm^3^ or ulcerated tumors were euthanized in accordance with the protocols approved by the Institutional Animal Care and Use Committees of Yale University.

### Immune cell isolation from implanted, lung and lymphoid organ

Subcutaneously implanted solid tumors were dissected, cut into small pieces and digested with 5mL Collagenase IV buffer (1x HEPES buffer, 0.5mg/mL Collagenase IV, 20μg/mL DNase in 1x HBSS with MgCl_2_ and CaCl_2_) for 30-40 minutes at 37°C. Samples were then run on the default lung_02 program on a gentleMACS Dissociator instrument (Miltenyi Biotec). Digestion was neutralized by adding 5mL 1% HI-FBS RPMI-1640. Tumor samples were filtered with 70 μm cell strainers and then washed with 1% HI-FBS RPMI-1640. Cell suspensions were then processed with Mouse CD45 (TIL) MicroBeads (Miltenyi Biotec) to enrich for CD45^+^ TILs. For autochthonous lung tumors, mice were injected intravenously with CD45-PE-CF594 (1:100, clone: 30-F11), 2-3 minutes before sacrifice as described previously (Joshi et al., 2015). Lungs were harvested, run on lung_01 program with gentleMACS Dissociator and digested with Collagenase IV buffer as described above, for 30-40 minutes at 37°C. Samples were then run on the lung_02 program, neutralized by 1% HI-FBS RPMI-1640, filtered with 70 μm cell strainers, washed with 1% HI-FBS RPMI-1640 and treated with RBC lysis buffer (eBioscience) to remove red blood cells. Tumor-draining lymph nodes and spleens were harvested and processed as described previously (Damo et al., 2020).

### Flow cytometry

Single cell suspensions were stained with antibodies for surface markers. For I-Ab/GP_66-77_-specific tetramer (NIH Tetramer Core Facility) staining, cells were incubated 2-3 hours at 37°C prior to surface staining. For intracellular staining, cells were processed using either Foxp3/Transcription Factor Staining Buffer Set (eBioscience) for transcription factors, or Fixation/Permeabilization Solution Kit (BD Cytofix/Cytoperm) for cytokines. Cells were washed and resuspended in FACS Buffers (PBS + 0.5% HI-FBS) until data collection. Flow cytometry was performed with LSR II flow cytometer (BD Bioscience), and analyzed by FlowJo software (version 10, TreeStar).

### CIBERSORT analysis

Fractions of different cell types were estimated with CIBERSORT (Newman et al., 2015), a cell type deconvolution program that derives cell type-specific abundances from bulk RNA-seq data. Here bulk RNA-seq data were downloaded from the whole TCGA LUAD cohort. Among the 594 samples with FPKM data, the first sample was used for each series of primary tumor samples. In total, 513 samples of primary tumor from different cases were selected for deconvolution. Reference marker gene expression profile (GEP) matrix was assigned with a widely used leukocyte signature matrix LM22 (Newman et al., 2015) provided by CIBERSORT program. Quantile normalization was disabled, and 100 permutations were used as default to calculate P values. From 22 cell subsets of CIBERSORT output using LM22 matrix, “B cells naïve”, “B cells memory” and “plasma cells” were combined as “B cell lineage”; “T cells CD4 memory resting” and “T cells CD4 memory activated” were combined as “CD4 memory T cells”; “NK cells resting” and “NK cells activated” were combined as “NK cells”; “macrophages M0”, “macrophages M1”, and “macrophages M2” were combined as “macrophages”; “dendritic cells resting” and “dendritic cells activated” were combined as “dendritic cells”; “mast cells resting” and “mast cells activated” were combined as “mast cells”.

### Single-cell RNA sequencing analysis

Data was downloaded from GEO with accession code GSE131907 (Kim et al., 2020). The raw UMI count matrix and annotations were used. Data normalization and analysis were performed with R package Seurat (version 3.1.0) and only cells from lungs or lymph nodes were used with original annotations “nLung”, “tLung”, “nLN”, or “mLN”. For the analysis of CD4^+^ T cells and B cells, 49,901 cells with original annotation “T/NK cells” and 22,592 cells with original annotations “B lymphocytes” were selected for the following analysis. UMI counts were log-normalized using function *NormalizeData* with parameter *normalization.method* set to “LogNormalize”. PCA was run based on the normalized data and clusters were called with *FindNeighbors* and *FindClusters* using the top 20 PCs and resolution set to 0.5 or 0.25 for T/NK cells or B cells, respectively. For both T/NK cells and B cells, dimension reduction and visualization were performed with UMAP using the top 15 PCs. Relative abundances for marker genes were calculated as Z-scaled average of log2(RC+1), where RC is relative counts that is calculated with *NormalizeData* and *normalization.method* set to “RC”.

### Survival analysis and correlation analysis

Survival analysis and pairwise correlation analysis of gene expression signatures were performed using web server GEPIA2 (Tang et al., 2019), based on TCGA and GTEx databases. 478 LUAD tumor samples were included in survival analysis. For each signature gene set, LUAD cohort was divided into high and low expression groups by median value (50% cutoff). The signature gene sets were defined as following based on previous studies: GC B cell: *AICDA, BATF, BACH2, BCL6, CD79A, CD79B, CD86, DOCK8, IRF4, IRF8, MYC* (Cabrita et al., 2020; Milpied et al., 2018). TFH: *ASCL2, BCL6, CD4, CD200, CXCR5, IL4, IL21, IL6ST, MAF, PDCD1, SH2D1A, TOX2* (Cillo et al., 2020; Vella et al., 2019). Th1: *BHLHE40, CXCR3, CD4, IFNG, IL2, IL12RB2, STAT4, TBX21*. CD8 effector cell: *CD8A, CD8B, CX3CR1, GZMA, GZMB, KLRG1, PRF1, ZEB2*. Th17: *CD4, IL6R, IL17A, IL17F, IL23R, RORA, RORC, STAT3*. NK cell: *CD160, CD244, GNLY, GZMB, KLRC3, KLRF1, NKG2A, NKG7*. Survival analyses were performed with log-rank Mantel-Cox test. Pairwise correlation analyses of gene expression signature were performed with two-tailed Pearson correlation test.

## Acknowledgements

This work was supported by grants from the NCI K22CA200912 (N.S.J.), NCI 1RO1CA237037-01A1 (N.S.J.), NIH R37AR40072 (J.C.), NIH AR074545 (J.C.), Yale SPORE in Lung Cancer 1P50CA196530 (N.S.J. and J.C.), and a Pilot Grant from Yale Cancer Center (N.S.J. and J.C.). C.C. was supported by the Gruber Science Fellowship. We thank N.S.J. and J.C. laboratory members for reviewing the manuscript and T. Mao, Z. Chen, C. Yang, and N. Ruddle for helpful discussions. We also thank the Yale Flow Cytometry Core, Yale School of Medicine Histology Facility and NIH Tetramer Core Facility. SW_HEL_ mice were kindly provided by Dr. Robert Brink. Figures 2A, 4D, 7B, S4D, S7A and graphical abstract were created with BioRender.com.

## Author contributions

C.C., J.C. and N.S.J. designed the study and wrote the manuscript. C.C., P.C., and E.F. performed experiments and generated primary data. C.C. and J.W. performed computational analysis of human data. P.C., K.A.C., M.D., S.C., S.C.E. and H.Z. provided helpful insight and contributed key reagents. J.C. and N.S.J. supervised data analysis and experiments.

## Declaration of interests

The authors declare no competing interests.

## Supplemental Information

**Figure S1.**
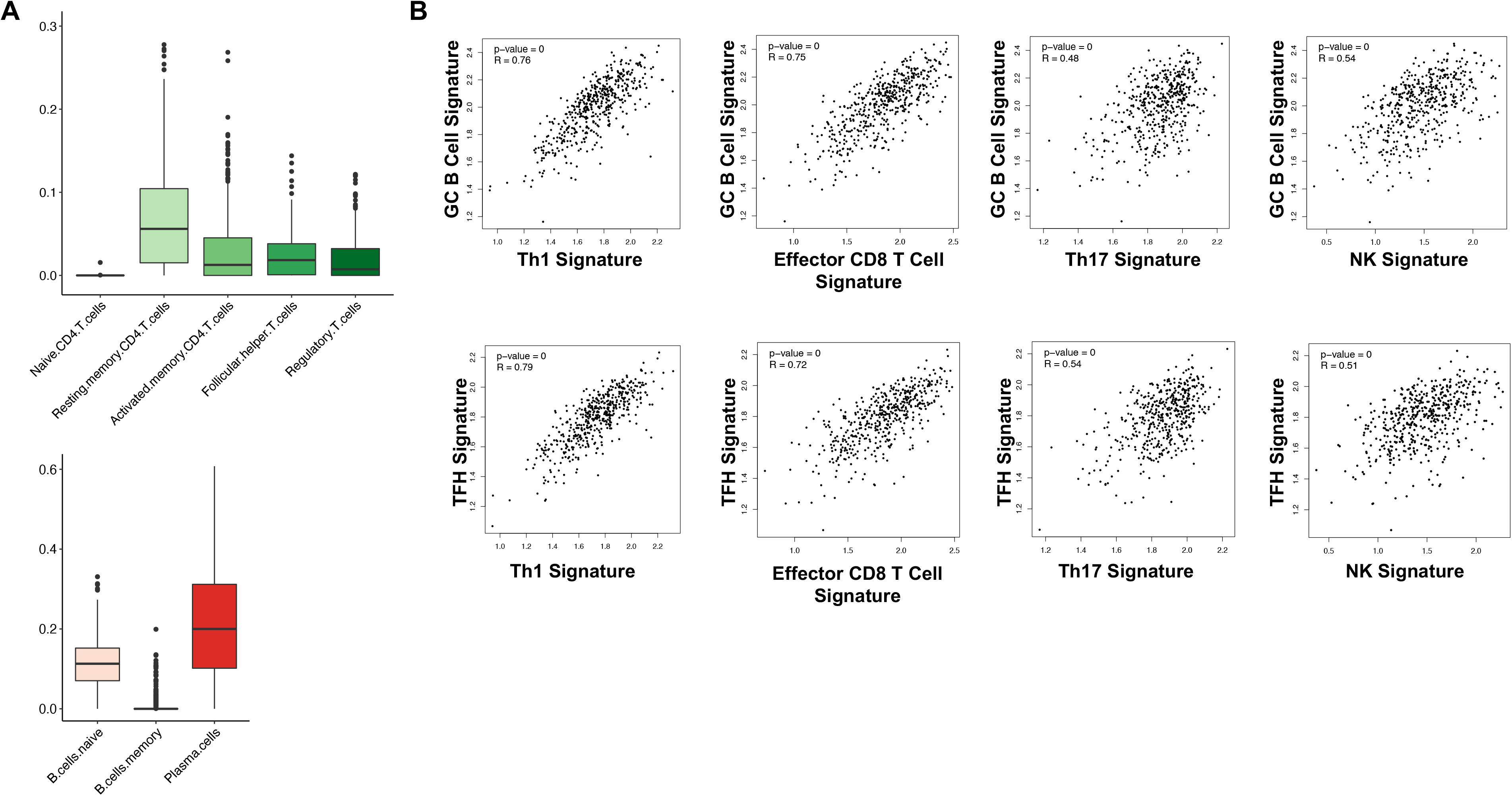
Analyses of T and B cells in LUAD samples from TCGA. **A,** Estimated fractions of subtypes in CD4^+^ T cells and B cell lineage described in **Figure 1A**, from CIBERSORT analysis of TCGA-LUAD samples (n=513). **B,** Correlation analyses among expression signatures of GC B cells, TFH, Th1, CD8 effector cells, Th17, NK cells in lung adenocarcinoma patients. Two-tailed Pearson correlation test was performed on expression signatures of TCGA-LUAD cohort (n=478).

**Figure S2.**
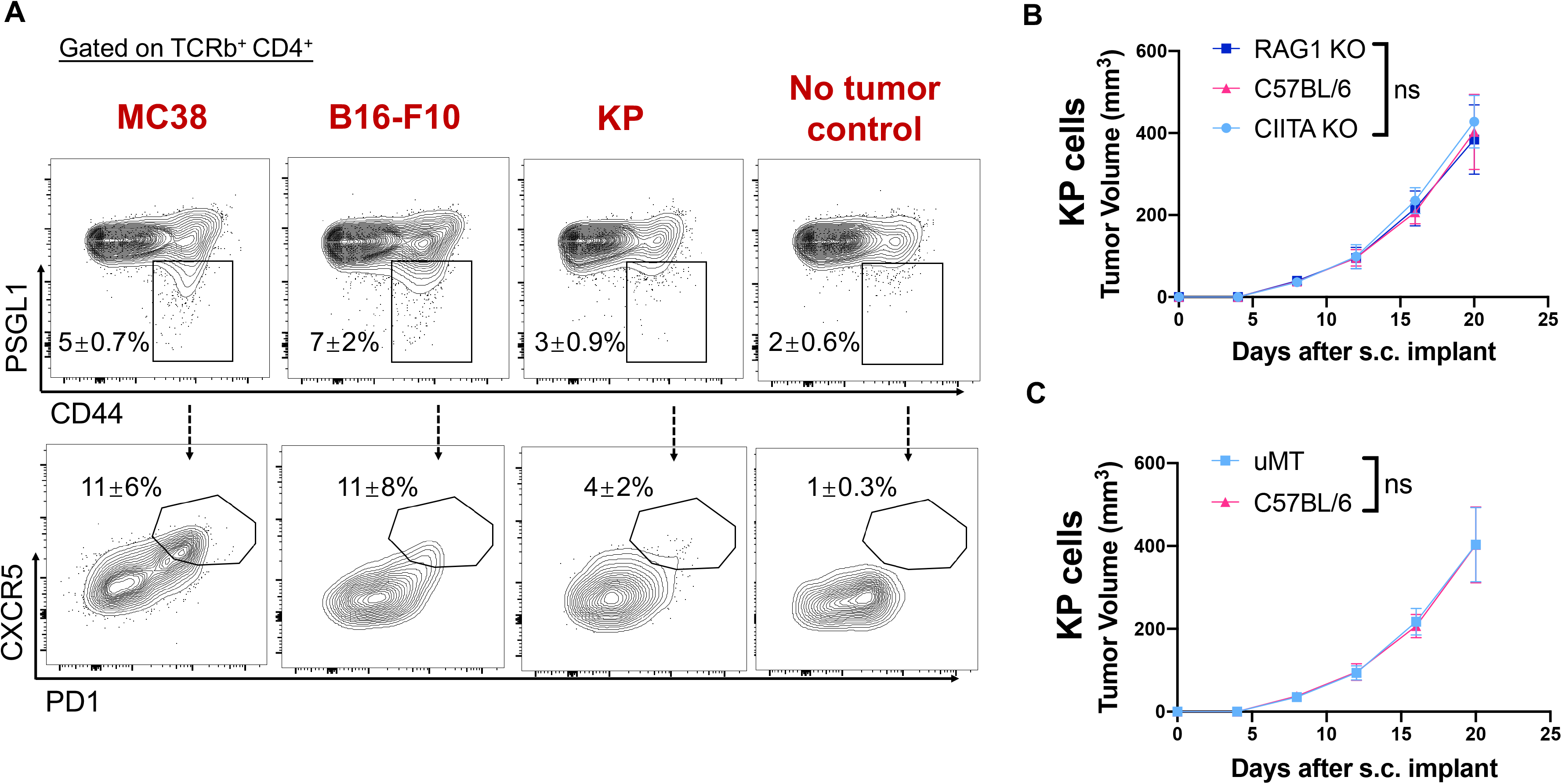
TFH cell responses in murine tumor models. **A,** Representative flow plots showing TFH gating in draining LNs of tumor-bearing C57BL/6 mice (pre-gated on TCRb^+^ CD4^+^). 5 × 10^5^ MC38, B16-F10 or KP tumor cells were implanted subcutaneously. Flow cytometric analyses were performed on day 10 – 12. Data (mean ± SD) represented 4 – 5 mice/group. **B – C,** Tumor growth curves: C57BL/6, RAG1 KO, CIITA KO (**B**), and uMT (**C**) mice were implanted subcutaneously with 2 × 10^5^ KP. Study end points: tumor size >1cm^3^, or signs of ulceration, infection, bleeding. Data (mean ± SD) were pooled, from 3 – 10 mice/group/experiment, and representative of at least 2 independent experiments. Two-way ANOVA. *P < 0.05, **P < 0.01, ***P < 0.001, ****P < 0.0001.

**Figure S3.**
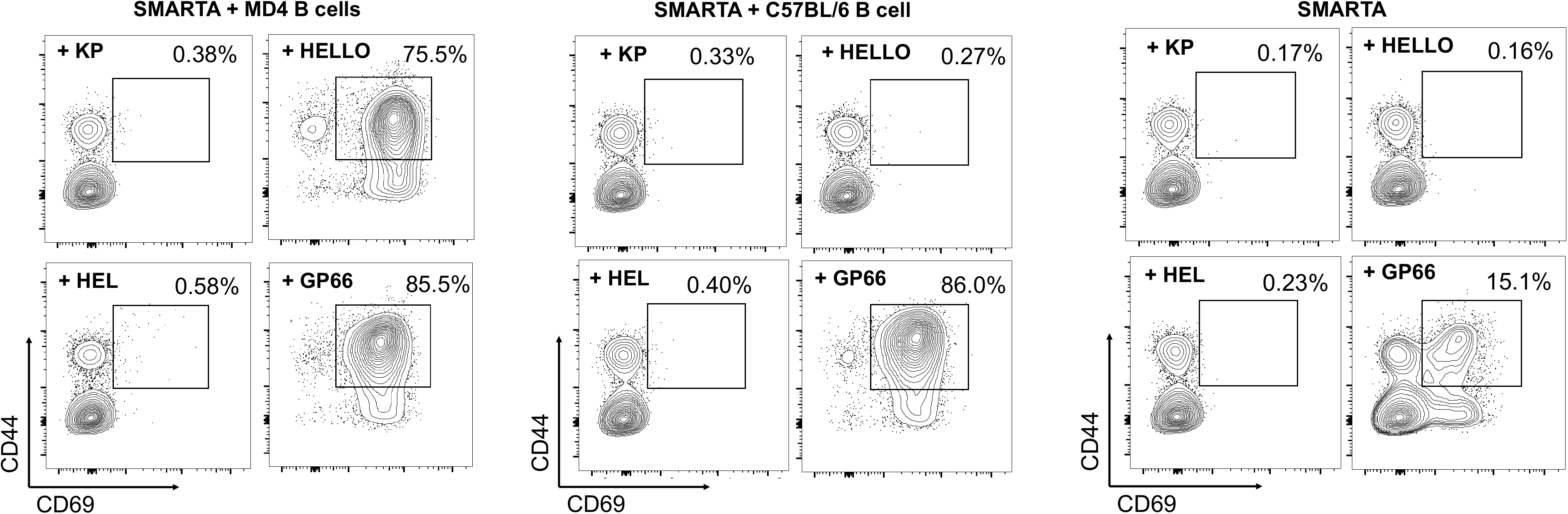
KP-HELLO lung adenocarcinoma cell line induces antigen-specific CD4^+^ T cell responses in vitro, via help from antigen-specific B cells. Representative flow plots showing the expression of CD69 and CD44 on SMARTA CD4^+^ T cells, 48 hours after *in vitro* co-culture with MD4 B cells or C57BL/6 B cells, and with the supernatant of KP, KP-HELLO, 500ng/mL HEL or 5ug/mL GP_61-80._

**Figure S4.**
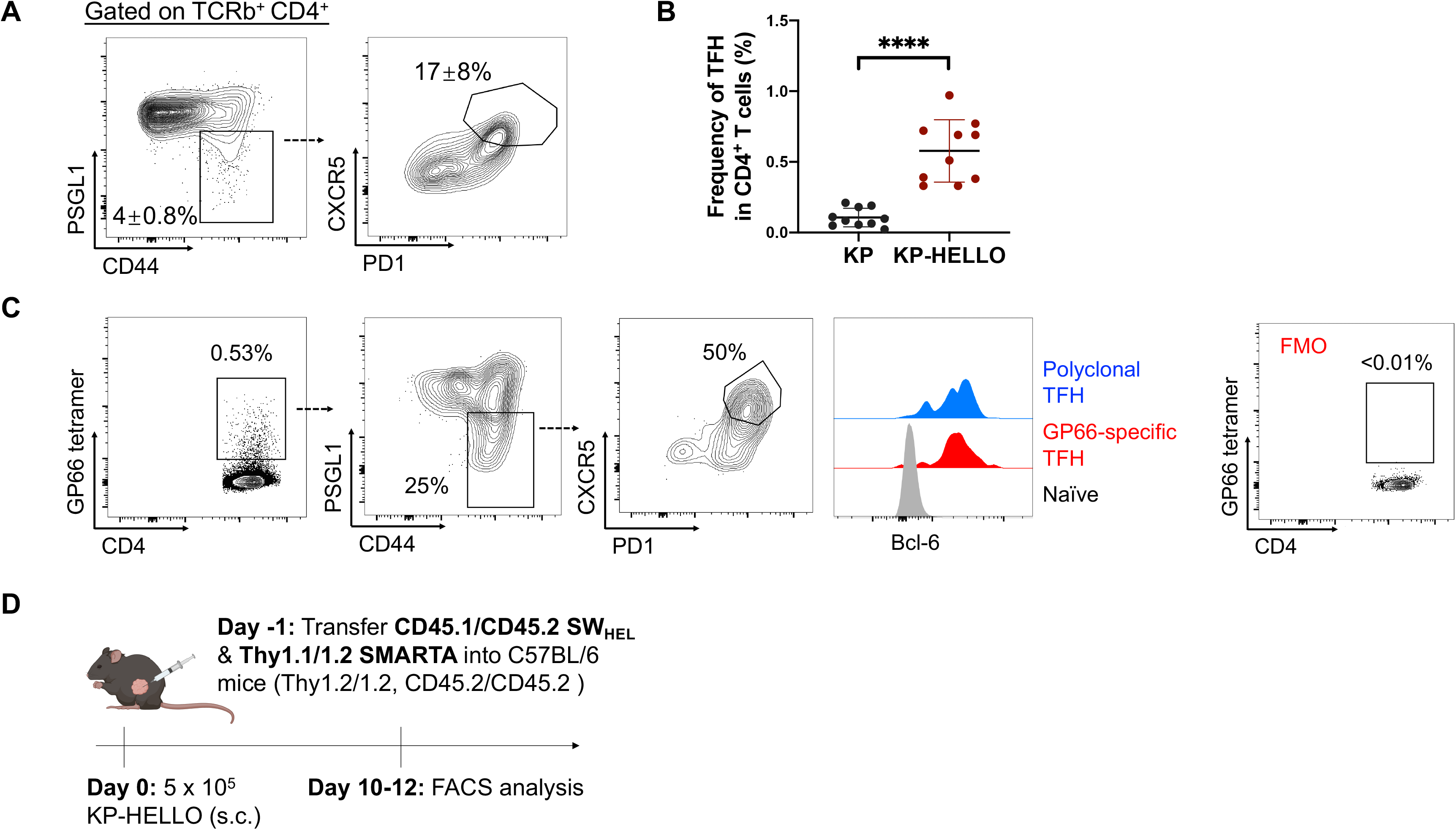
KP-HELLO lung adenocarcinoma elicits antigen-specific TFH and GC B cell responses in vivo. **A – B**, Representative flow plots (**A**) and frequency (**B**) of polyclonal CD4^+^ TFH cells from draining LNs of KP-HELLO tumor-bearing C57BL/6 mice. Data (mean ± SD) were pooled, from 3 – 6 mice/group/experiment, and representative of 3 independent experiments. Two-tailed Student’s t-test. *P < 0.05, **P < 0.01, ***P < 0.001, ****P < 0.0001. **C,** Representative flow plots of I-A^b^/GP_66-77_-specific CD4^+^ TFH cells from draining LNs of KP-HELLO tumor-bearing C57BL/6 mice. **D,** Experimental design of **Figure 2D – 2E**.

**Figure S5.**
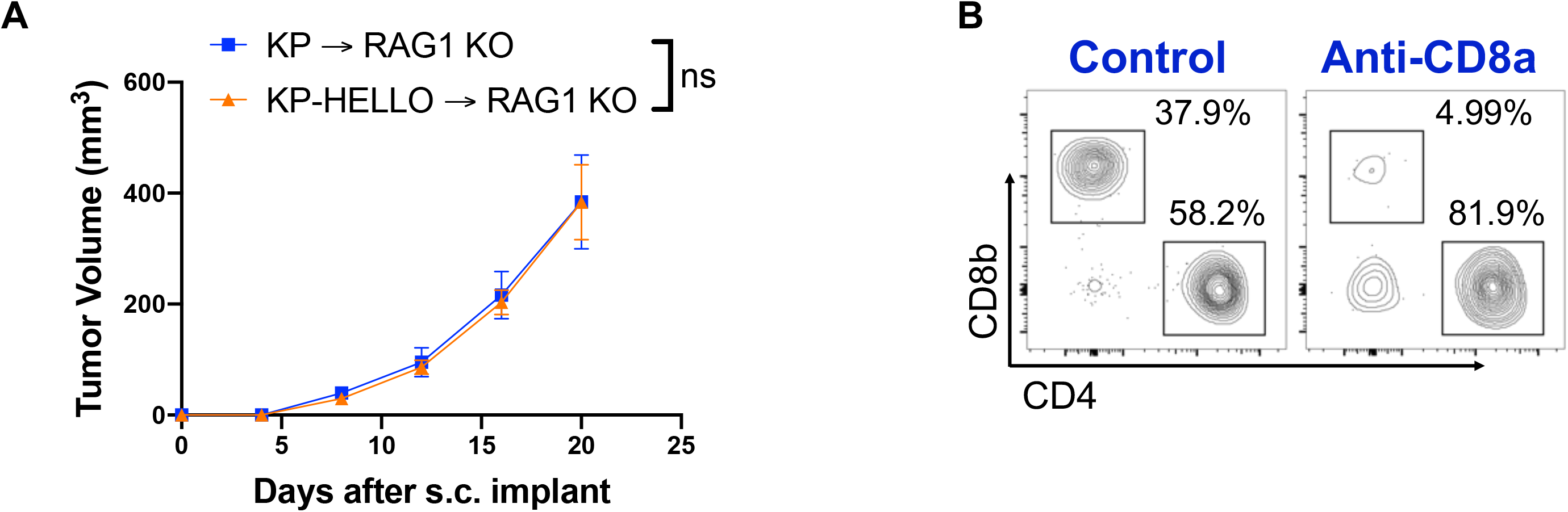
T and B cells are necessary for optimal tumor control for KP-HELLO tumors. **A,** RAG1 KO mice were implanted subcutaneously with 2 × 10^5^ KP or KP-HELLO. Study end points: tumor size >1cm^3^, or signs of ulceration, infection, bleeding. Data (mean ± SD) were pooled, from 3 – 10 mice/group/experiment, and representative of 2 independent experiments. Two-way ANOVA. *P < 0.05, **P < 0.01, ***P < 0.001, ****P < 0.0001. **B,** Representative flow plots showing T cells in peripheral blood samples from control C57BL/6 and anti-CD8a treated mice, 10 days after the initiation of antibody administration.

**Figure S6.**
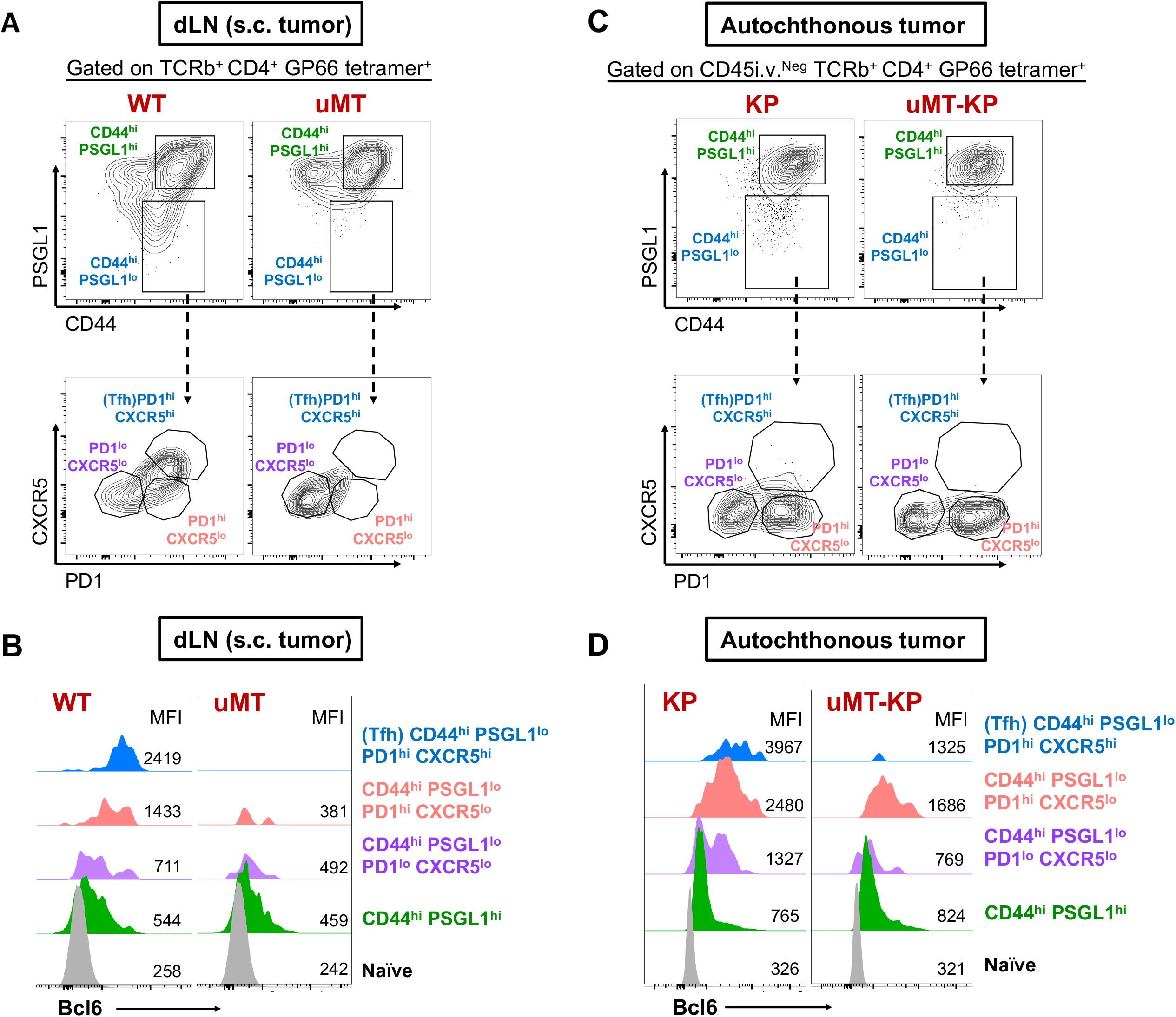
B cells are required to drive effective antigen-specific TFH response. **A and C,** Gating strategy of I-A^b^/GP_66-77_-specific CD4^+^ T cell subsets in draining LNs of subcutaneously (s.c.) implanted KP-HELLO tumors (**A**) or tumor lung tissues of autochthonous KP-HELLO tumors (**C**). **B and D,** Histogram displaying Bcl6 expression in naïve, I-A^b^/GP_66-77_-specific CD44^hi^ PSGL1^hi^, CD44^hi^ PSGL1^lo^ PD1^lo^ CXCR5^lo^, CD44^hi^ PSGL1^lo^ PD1^hi^ CXCR5^lo^ and TFH cells (CD44^hi^ PSGL1^lo^ PD1^hi^ CXCR5^hi^) from dLN of subcutaneously (s.c.) implanted KP-HELLO tumors (**B**) or tumor lung tissues of autochthonous KP-HELLO tumors (**D**). Data were representative of 3 independent experiments.

**Figure S7.**
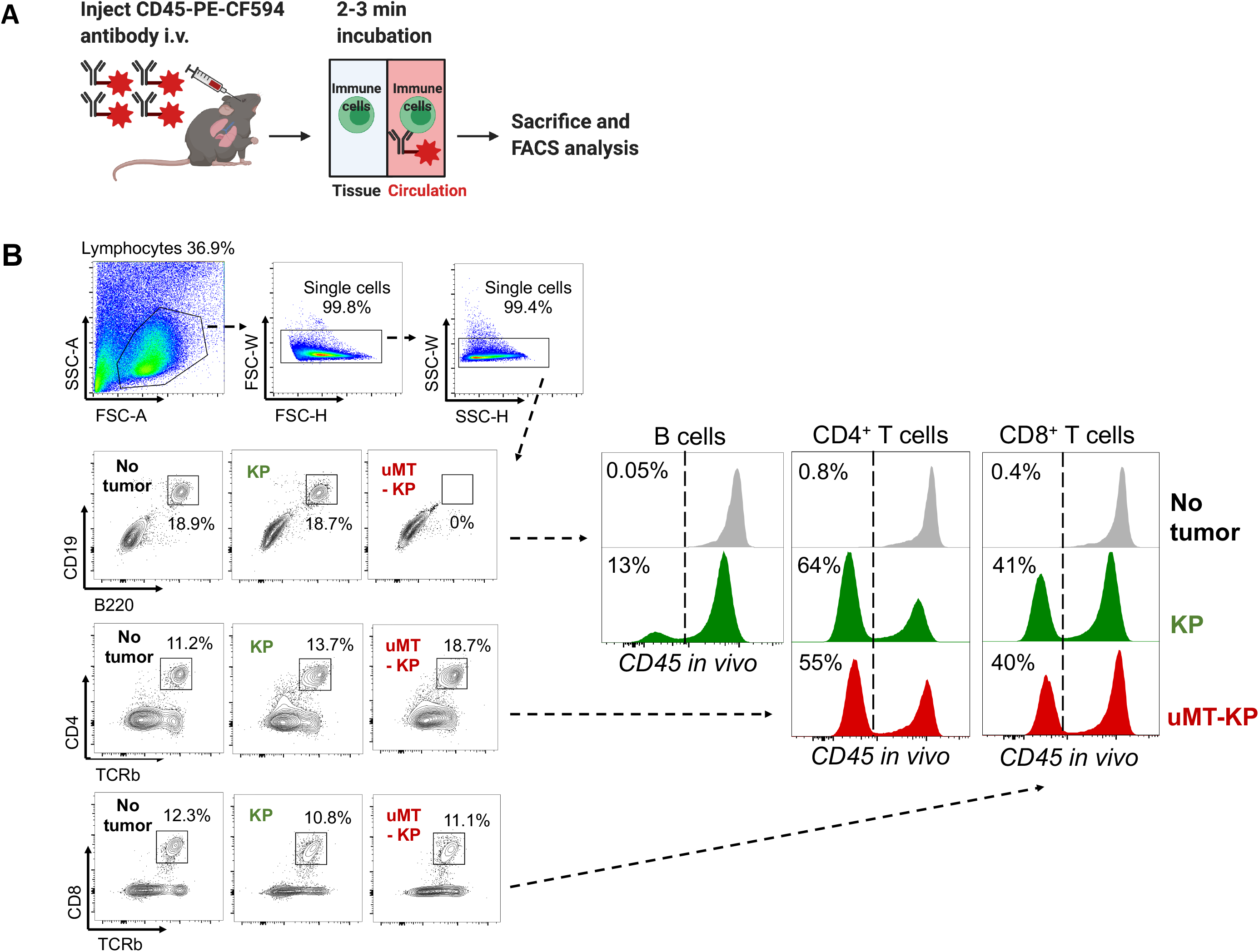
Experimental strategy for flow cytometric analysis of tumor-infiltrating immune cells in lung tissue. **A,** Diagram showing the technique for labeling circulating immune cells in lung for FACS analyses. 2-3 minutes prior to sacrifice, PE-CF594-conjugated CD45 antibodies were injected intravenously. Immune cells in circulation were labelled, while the time was not enough for antibodies to diffuse into tissue and bind to tumor-infiltrating immune cells. **B,** Gating strategy and histogram of CD45-PE-CF594 (*in vivo* labeling) showing tissue-infiltrating B cells, CD4^+^ T cells and CD8^+^ T cells in non-tumor-bearing mice, or tumor-bearing KP and uMT-KP mice. KP or uMT-KP mice were intratracheally infected with 5 × 10^4^ PFU Lenti-HELLO-Cre. Tumor lung tissues and LNs were harvested at 8 weeks post infection.

**Figure S8.**
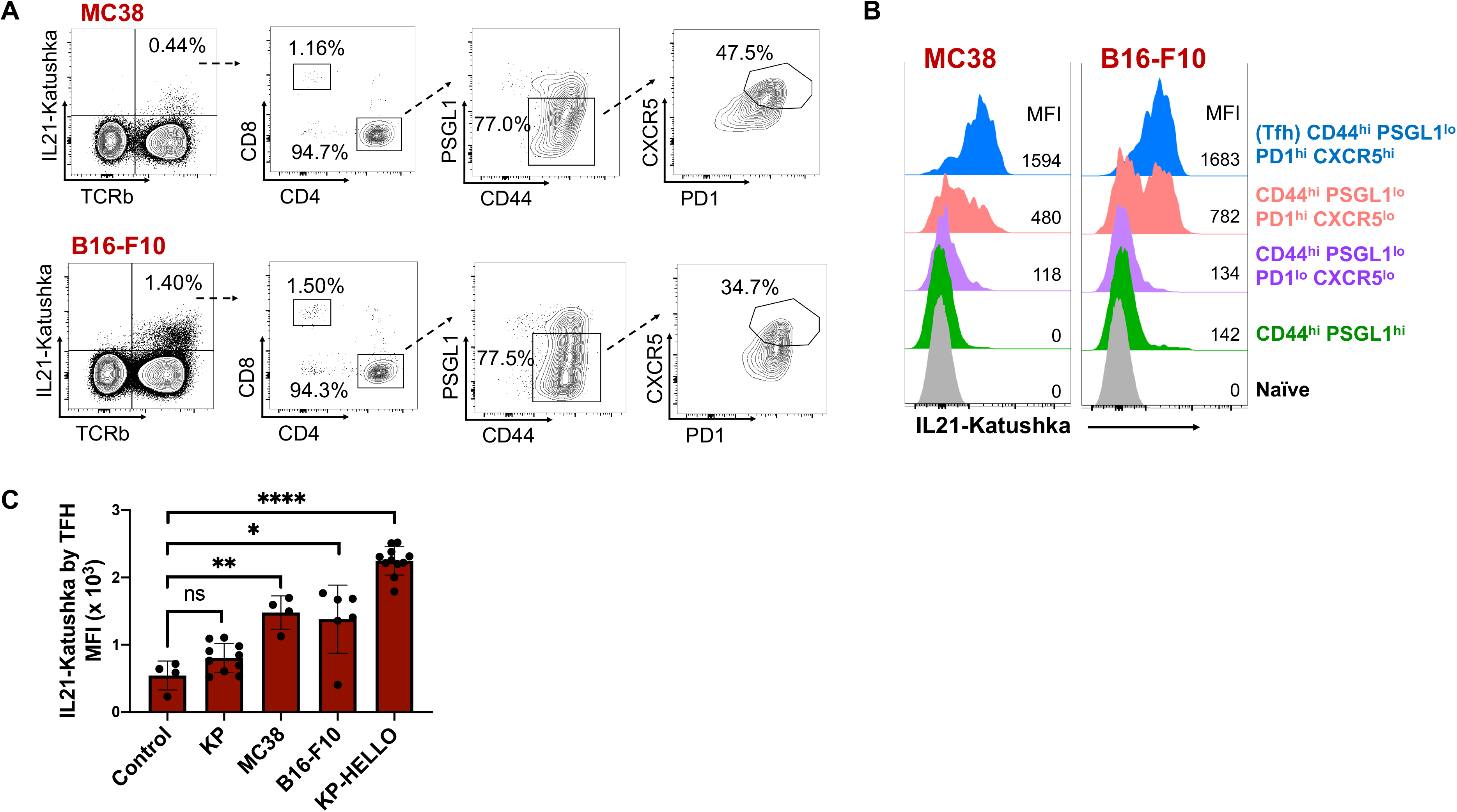
TFH-derived IL-21 in murine tumor models. **A – B,** Representative flow plots showing IL21-expressing TFH cells (**A**) and histogram displaying the expression of IL21 through different stages of TFH differentiation (**B**) in draining LNs of MC38 and B16-F10 tumor-bearing mice. Gating strategy was shown in **Figure S6**. Flow cytometric analyses were performed on day 10 – 12. **C,** Expression of IL21 by TFH cells (CD44^hi^ PSGL1^lo^ PD1^hi^ CXCR5^hi^) in draining LNs of KP, MC38, B16-F10 and KP-HELLO tumors. Data (mean ± SD) were pooled, from 3 – 6 mice/group, and representative of 2 independent experiments. Two-tailed Student’s t-test. *P < 0.05, **P < 0.01, ***P < 0.001, ****P < 0.0001.

## Notes

### Competing Interest Statement

The authors have declared no competing interest.

